# North and East African mitochondrial genetic variation needs further characterization towards precision medicine

**DOI:** 10.1101/2021.12.10.472079

**Authors:** Anke Fähnrich, Isabel Stephan, Misa Hirose, Franziska Haarich, Mosab Ali Awadelkareem, Saleh Ibrahim, Hauke Busch, Inken Wohlers

**Author notes:** Equal contributing first authors (Anke Fähnrich, Isabel Stephan). Equal contributing last authors (Hauke Busch, Inken Wohlers).

## Abstract

**Introduction:** Mitochondria are maternally inherited cell organelles with their own genome, and perform various functions in eukaryotic cells such as energy production and cellular homeostasis. Due to their inheritance and manifold biological roles in health and disease, mitochondrial genetics serves a dual purpose of tracing the history as well as disease susceptibility of human populations across the globe. This work requires a comprehensive catalogue of commonly observed genetic variations in the mitochondrial DNAs for all regions throughout the world. So far, however, certain regions, such as North and East Africa have been understudied.

**Objectives:** To address this shortcoming, we have created the most comprehensive quality-controlled North and East African mitochondrial dataset to date and use it for characterizing mitochondrial genetic variation in this region.

**Methods:** We compiled 11 published cohorts with novel data for mitochondrial genomes from 159 Sudanese individuals. We combined these 641 mitochondrial sequences with sequences from the 1000 Genomes (n=2,504) and the Human Genome Diversity Project (n=828) and used the tool haplocheck for extensive quality control and detection of in-sample contamination, as well as Nanopore long read sequencing for haplogroup validation of 18 samples.

**Results:** Using a subset of high-coverage mitochondrial sequences, we predict 15 potentially novel haplogroups in North and East African subjects and observe likely phylogenetic deviations from the established PhyloTree reference for haplogroups L0a1 and L2a1.

**Conclusion:** Our findings demonstrate common hitherto unexplored variants in mitochondrial genomes of North and East Africa that lead to novel phylogenetic relationships between haplogroups present in these regions. These observations call for further in-depth population genetic studies in that region to enable the prospective use of mitochondrial genetic variation for precision medicine.

## Introduction

Mitochondria are energy-producing, double-membrane-bound organelles that contain their own genome and that are central to many processes in eukaryotic cells (1). The mitochondrial (MT) genome is circular and typically 16,569 bases long in humans (2). Mitochondrial dysfunction is often linked to inefficient oxidative phosphorylation (OXPHOS) capacity with compromised ATP and increased mitochondrial reactive oxygen species (ROS) production that change cellular function and signaling (3). Dysfunction accompanies various pathologic conditions, such as metabolic diseases (4), neurodegenerative disorders (5), and autoimmune conditions as well as cancer (6) and is also linked to aging (7). Its root cause might stem from variations of the MT genome, which has thus moved into the center of research as a determining factor for complex diseases and precision medicine.

Since mitochondria are exclusively maternally inherited in humans, the mitochondrial DNA (mtDNA) phylogeny represents a global maternal genealogy, which has been summarized in an established and widely applied reference phylogenetic tree called PhyloTree (8,9). Its current version 17 is based on more than 24,000 mitochondrial sequences and constitutes more than 5,400 haplogroups, i.e., recurrent genetic variant profiles common in human mitochondrial DNA.

Recent studies assessed mitochondrial haplogroups in worldwide human diversity reference data sets from the 1000 Genome Project (1000G; n=2,504) (10,11), the Human Genome Diversity Project (HGDP; n=623 in (12); n=102 in (13)) and very recently gnomAD (14). Such mitochondrial phylogenetic studies have traced modern human settlement of the world (15) and support the hypothesis of Africa as origin of modern humans. The “Out-of-Africa” theory proposes the so-called “mitochondrial Eve”, which represents the mitochondrial Reconstructed Sapiens Reference Sequence (RSRS) mtDNA as the phylogenetic root and modern human origin (16).

Previous studies on mtDNA genetic variation in North and East African individuals largely focused on identifying and characterizing phylogenetic clades with respect to time of origin and geographic dispersal. However, with advances in next-generation sequencing technologies and decreasing sequencing costs, the wide-spread use of mitochondrial genetic variation for medical purposes becomes feasible. Many applications in this context can be considered as precision medicine, i.e. they provide personalized diagnostics or treatment. They fall into two areas: primary mitochondrial diseases and common diseases (17). Primary mitochondrial diseases are caused by a genetic defect that impairs MT function, resulting into a range of severe symptoms. They affect one of 4,300 people (18). To clinically diagnose these genetic diseases, the underlying causal variant needs to be identified on an individual basis, a process in which variants that occur in healthy individuals are ruled out. Thus, reference data that is representative of human genetic mitochondrial variation will improve diagnosis of primary mitochondrial diseases. The second area of application are common, polygenic diseases; differences in mitochondrial DNA have been associated with a range of them. Considering the central biological role of mitochondria, mtDNA variation is particularly promising for disease stratification, identification of disease modifiers and inclusion into personalized risk assessment, e.g. via polygenic risk scores. MtDNA for precision medicine in common diseases is a comparably novel field with large potential (17). Owing to these recent developments, we here view mitochondrial genetic variation of North and East Africa from the point of its application in primary mitochondrial disease diagnostics and prospective precision medicine.

In this work, we address the state of mtDNA reference data from the regions of North Africa and East Africa. As of now, there are limited genetic data available from this world region, as it is not included in large genome projects such as 1000G (19), gnomAD (20) or Topmed (21). An exception is Egypt, for which an initial reference dataset has been compiled (22) and efforts are ongoing to construct a much larger Egyptian genome reference towards genomics-based precision medicine (23).

To address this reference data limitation, we overall compiled and subsequently quality controlled data from overall 844 mitochondrial genomes (cf. Suppl. Table 1). These are from a newly sequenced Sudanese cohort (n=159), an Egyptian cohort with targeted mitochondrial sequencing (n=217) and an Egyptian cohort having whole genome sequencing data (n=110). Both Egyptian cohorts have previously been investigated for haplogroup distribution (22,24) and are here combined with mitochondrial sequencing data of ancient Egyptian mummies (n=90) (25) as well as mtDNA sequences from Genbank from individuals with origins in North or East African countries (n=268).

**Table 1:** Haplogroups having potentially novel and previously undescribed sub-haplogroups. The table columns denote, in this order, haplogroup, recurrent novel variants, the number of samples after strict filtering (# HQ; out of n=146) and after lenient filtering (#LQ; out of n=641-146=495) and the Soares score (48), which represents the previous occurrence of the variant in worldwide data sets.

| Haplogroup | Subhaplogroup variant | # HQ | # LQ | Soares score |
| --- | --- | --- | --- | --- |
| HV1b+152 | 8640T; 4590G | 2 | 1 | 0; 0 |
| I | 567C | 3 | 0 | 0 |
| I | 567C; 4772C | 2 | 3 | 0; 2 |
| L0a1a1 | 8869G | 2 | 1 | 1 |
| L0a1a1 | 8017T | 3 | 9 | 0 |
| L2a1c | 5186G; 9116C | 2 | 0 | 2; 2 |
| L2a1+143 | 7897A; 398C; 9389G | 2 | 0 | 1; 2; 2 |
| L2e1 | 13356C | 2 | 0 | 0 |
| L3d5a | 15457A; 14410A | 2 | 1 | 0; 1 |
| L3e4a | 470G; 9455G | 2 | 2 | 0; 1 |
| L3e5 | 711C | 2 | 2 | 0 |
| L3f1a1 | 12441C; 736T; 328G; 15151G; 13967T; 14554G | 3 | 3 | 0; 0; 0; 1; 2; 2 |
| M1a2 | 16182G; 2858G; 15401G; 1282A | 2 | 0 | 1; 1; 1; 1 |
| R0a2c | 5981C | 2 | 1 | 0 |
| V | 9126C | 2 | 1 | 0 |

In addition to providing additional mitochondrial genomes from this understudied region, we reanalyzed all available next generation sequencing data using a novel tool, called haplocheck (26), to identify potential sample contamination and phantom variants (27). Haplocheck is a recent tool which allows to detect sample contaminations from next-generation sequencing data by assessing if multiallelic variant calls can be broken down into a mixture of two haplogroups, one denoted as “major”, the other as “minor”. Presence of two haplogroups indicates that genetic material of two individuals was jointly sequenced and thus DNA from the individual relating to the major haplogroup contaminated with DNA from an individual carrying the minor haplogroup. Our combined and new quality-controlled data results in 146 exceptionally high-quality and 641 high quality mitochondrial genome sequences after quality control that we used to phylogenetically re-characterize genetic MT variation in North and East Africa with respect to worldwide mtDNA diversity. We find that the haplogroups of the region cover nearly all worldwide clades and identified 15 potential novel subhaplogroups, among them two that descend from L0a1a1 and and two that are related to L2a1, for which sequences from North and East Africa suggest a different phylogenetic relationship as described by the reference PhyloTree.

## Results & Discussion

### North and East African mitochondrial data collection and filtering

To comprehensively cover the phylogenetic tree of the North and East African population we sequenced a cohort of 159 Sudanese individuals which we combined with mitochondrial sequencing data from 217 Egyptians. An additional 110 MT genomes were extracted from whole genome sequencing data of Egyptians from Pagani et al. (24) and Wohlers et al. (22). We further included mapped mitochondrial sequencing data from 90 ancient Egyptian mummies from Schuenemann et al. (25); for three of these mummies, WGS data were available (cf. Supplementary Table 1 for details).

As controls, we used two worldwide whole-genome references, 1000G with 2,504 samples (19) and HGDP with 828 samples (28) from which we generated mitochondrial DNA sequences. Further, we obtained quality-controlled and aligned Genbank sequences from countries in North and East Africa from the supplement information of McInerny et al. (29), which we manually traced to their eight respective publications provided in Suppl. Table 1. Sampling location or ethnicity as well as number of sequences are provided in Figure 1.

**Figure 1:**
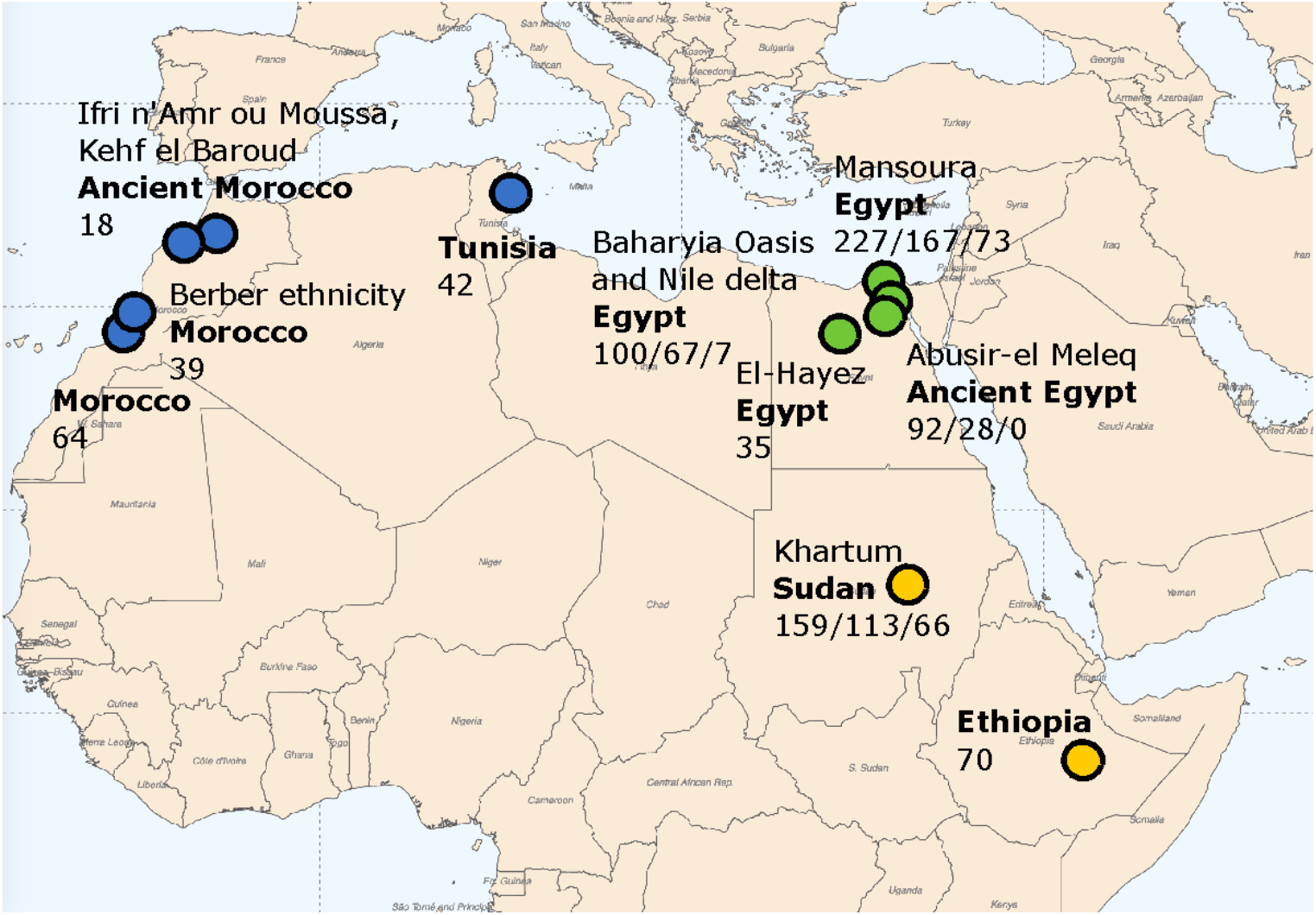
Mitochondrial DNA data compiled from North and East Africa. Blue color denotes cohorts referred to as North African, green color cohorts refer to Northeast African and yellow color cohorts refer to East African. Cohorts analyzed from raw next-generation sequencing data are denoted by three numbers: The overall number of samples; the number of samples remaining after lenient QC filtering; and the number of samples remaining after strict QC filtering. Locations with only one number refer to Genbank MT sequences obtained via McInerny et al. (29). Whenever Northeast Africa is not shown separately, its cohorts are included in North Africa and East Africa.

Five of eight studies relating to Genbank entries included only specific haplogroups. These are two studies on haplogroup U6 (Maca-Meyer et al. (30), n=11; Olivieri et al. (31), n=28) in Berbers from Morocco; two studies on haplogroup L3 in Morocco (Harich et al. (32), n=8) and Ethiopia (Soares et al. (33), n=70); and one study of both M1 and U6 in Moroccan (Pennarun et al. (34), n=56). We also included 18 Genbank MT sequences from ancient Moroccans (Fregel et al. (35)). Genbank-related studies that did not focus on a specific haplogroup covered 35 Egyptian (Kujanová et al. (36)) and 42 Tunisian mtDNA sequences (Costa et al. (37)).

To exclude study-specific batch effects and sequencing errors, we applied a two-step quality control (QC) filter on all next-generation sequencing-based North and East African data sets. This was important to correctly assess haplogroup distributions and identify putatively new haplogroups. For the filter step, we kept all samples from Genbank as well as those (i) having a haplocheck-reported major haplogroup equal to a minor haplogroup, and (ii) having contamination assessed and not detected (i.e. “NO”). An overall of 641 samples from four next-generation sequencing-based North and East African data sets and from eight Genbank studies passed this lenient filter (cf. Suppl. Table 11 and Suppl. Table 18 for aligned sequences). For the second QC filtering step, we excluded Genbank sequences, as contamination checks for them were unfeasible, and additionally required a minimum sequencing coverage greater than 1000x, which is considerably higher than the minimum 600x coverage required by haplocheck to detect a 1% or greater contamination (26). The number of total and filtered samples in each QC step are provided in Figure 1 for every cohort included. For mitochondrial sequences obtained from Genbank, only the overall number is provided.

### Haplogroup-based contamination assessment for diverse sequencing settings

We assessed potential contamination and related sequencing and variant calling metrics for all six sequencing-based datasets, including the worldwide references 1000G and HGDP (cf. Supplementary Tables 2-9 and 13-16 for details). Haplogroups assigned to 543 HGDP samples, for which Lippold *et al*. (12) determined haplogroups previously using targeted mitochondrial sequencing were largely consistent with the haplogroups assigned in this work (47% identical haplogroup; 86% a subclade of the previously assigned haplogroup; cf. Suppl. Table 17). Coverage of the 1000G and HGDP data sets is high with a median greater than 10,000X, while the other four cohorts have a significantly lower coverage of 840X (Egyptian MT sequencing), 670X (Egyptian WGS), 163X (ancient Egyptian mummies) and 992X for the Sudanese MT sequencing cohorts, respectively (see Figure 2 b).

**Figure 2:**
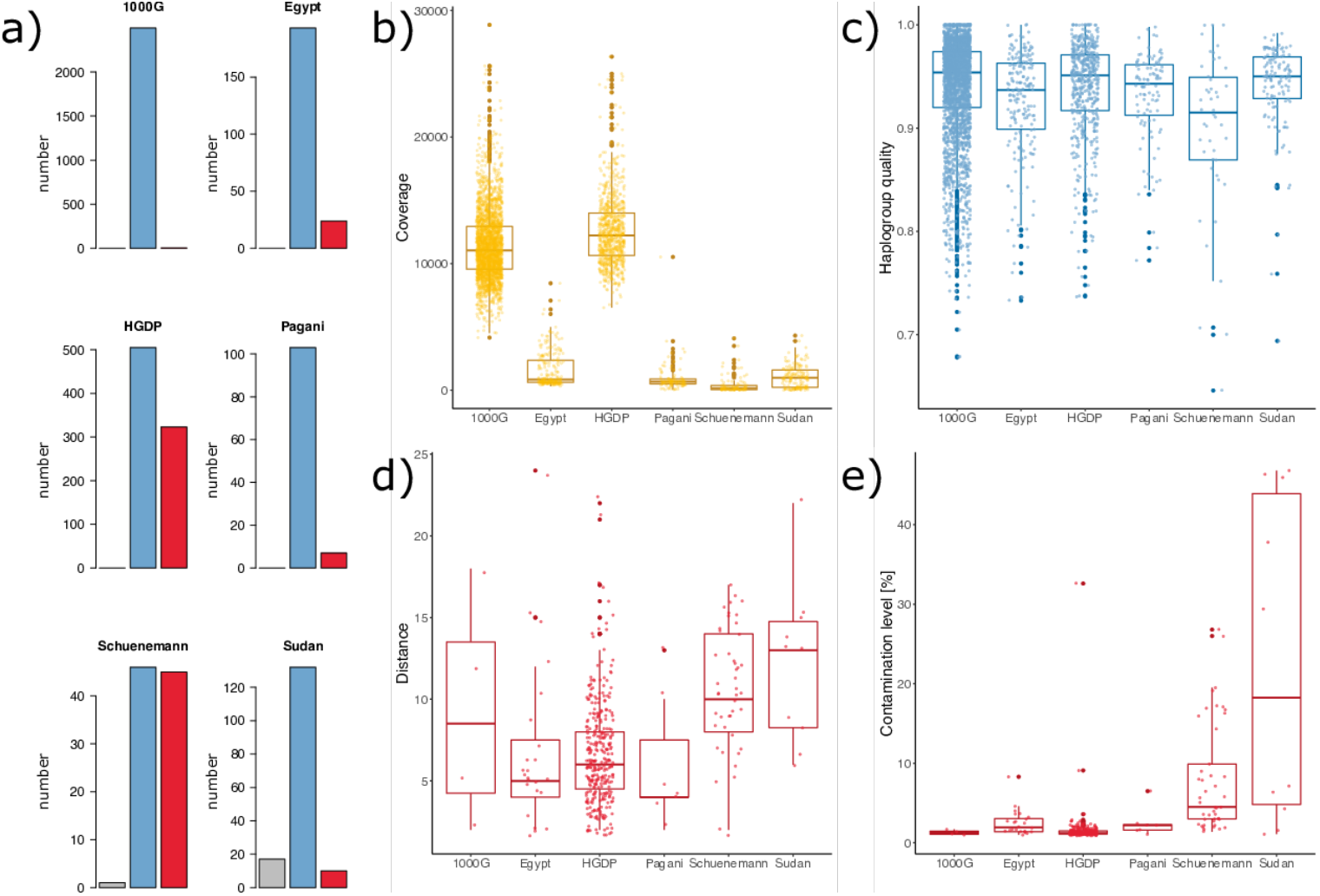
Characterization of the six sequencing-based data sets using numbers reported by haplocheck. (a) Histogram of number of samples with haplocheck contamination status not available (“NA”) (gray), not contaminated (“NO”) (blue) and predicted contaminated (“YES”) (red); (b) Box plots for sequencing coverage; (c) Box plots for haplogroup quality of those samples not predicted to be contaminated; (d) For those samples predicted to be contaminated, box plots of the distance between major and minor haplogroup, i.e. the number of phylogenetic nodes between them within the phylogenetic tree; (e) Box plots of the contamination levels for samples predicted as contaminated. Box plots display median and lower/upper quartiles; whiskers denote the most extreme data point no more than 1.5 times the interquartile range; outliers are data points extending beyond whiskers.

Contamination is identified using the computational tool haplocheck, which identifies polymorphic sites within samples. It attempts to decompose the variants into a so-called major and another, minor, haplogroup. If such decomposition is possible and coherent, a mix of a sample of the major haplogroup with another sample of the minor haplogroup is predicted and the percentage of minor haplogroup allele frequency reported as contamination level. Interestingly, we found a large heterogeneity in contamination calls among the studies. Few of the WGS-based 1000G samples are predicted to be contaminated, yet more than 30% of the WGS-based HGDP samples (Figure 2a), albeit at a low contamination level of 2% (median 1.2%, Figure 2e). Comparable, low level high distance contamination is predicted for WGS samples of Wohlers *et al*. and Pagani *et al*. Possibly, these contamination calls are caused by NUclear MiTochondrial DNA sequences (NUMTs) causing mapping artefacts, although a core genome coverage of 30x, a typically large haplogroup quality (Figure 2c) and a large phylogenetic distance between major and minor haplogroup detectable based on unusual mutation patterns (Figure 2d) indicate that this is rather not the case. High contamination levels (e.g. levels of 50% or 30%) are observed for very few samples largely from the Sudan cohort and may indicate partial swaps of sample material. For the ancient Egyptian mummy data set, haplocheck repeatedly identifies minor haplogroups related to H2a2a, which we consider an artifact caused by a lack of sequencing coverage that causes the revised Reference Sequence (rCRS) reference bases to be called, which refer to haplogroup H2a2a1. This is supported by the very low coverage for this cohort.

### Haplogroup validation with Nanopore long-read sequencing

We performed Nanopore long-read sequencing to validate haplogroups obtained from whole-genome sequencing data as well as from targeted mitochondrial sequencing data. We used 10 Egyptian samples whole genome-sequenced at 30x (covering subhaplogroups of H, L, M, and T). For all of them, the major haplogroup was confirmed. Further, we performed Nanopore sequencing for eight Sudanese samples for which mitochondrial sequencing data were available. The major haplogroup for all of them was confirmed, with the exception of L2a1+143, which according to Nanopore sequencing was assigned to the closely related haplogroup L2a1’2’3’4, with no contamination predicted (Suppl. Table 19). Manual inspection shows that all four variants that distinguish between L2a1’2’3’4 and L2a1+143 are clearly detectable from Nanopore sequencing, but have a large fraction of reads carrying the reference allele (chrM:143 16%; chrM:12693 16%; chrM:15784 24%; chrM:16309 7%). Interestingly, we observe a large number of predicted contaminations (n=13/18; 72%) with subhaplogroups of H2a2a, which, again, indicates a reference bias caused by a sequencing or variant calling artefact. Nanopore sequencing has a comparably low base accuracy and base calling errors affecting the reference base may be preferentially reported by MT DNA callers which have been designed for short-read data that has higher base accuracy. As technologies, protocols and tools for long-read data develop quickly, constant benchmarking is needed and has recently shown the validity for Nanopore-based somatic MT variant calling (38).

### Heteroplasmic variant calls

Besides homoplastic variants, which are monoallelic, mtDNA variants can be heteroplasmic, i.e. mtDNA of more than one allele is present in a sample or tissue. Highly accurate detection of heteroplasmic variants, especially at low heteroplasmy levels, was not the scope of our study and needs dedicated sequencing and analysis. Nevertheless, we investigated the overall number of homoplasmic and heteroplasmic variant calls provided by haplocheck to characterize the next-generation sequencing data sets (see Suppl. Tables 3, 5, 7, 9, 14 and 16). For those samples not predicted to be contaminated (and thus not affected by artificial heteroplasmic calls) the results are displayed in Figure 3 a) and b).

**Figure 3:**
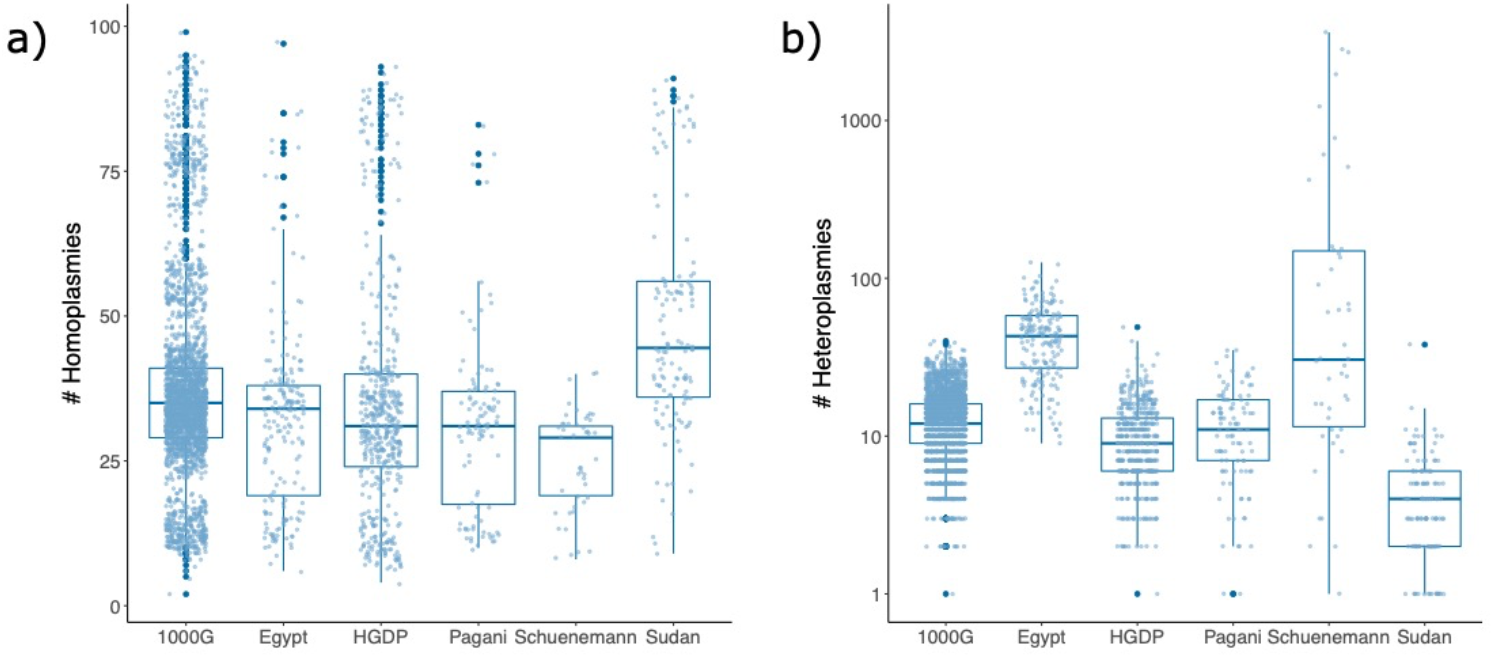
Boxplots of number of hompolasmic and heteroplasmic variant calls with respect to rCRS for all six next-generation sequencing-based data sets. Box plots display median and lower/upper quartiles; whiskers denote the most extreme data point no more than 1.5 times the interquartile range; outliers are data points extending beyond whiskers.

The number of homoplasmic variant calls reflects the haplogroups present in the respective data set, because the larger the haplogroup’s phylogenetic difference from the reference sequence rCRS haplogroup H2a2a1, the more variants will be called with respect to rCRS. As such, data sets containing for example many African haplogroups L will have more samples with many homoplasmic variant calls. This also explains the comparably large number of homoplasmic variant calls in the Sudanese data set. However, numbers of heteroplasmic variant calls are expected to be comparable between data sets and independent of the haplogroup content of the respective data set. This is however not the case and differences in the distributions of heteroplasmic variant calls can be observed between datasets. These differences indicate technical artefacts such as NUMTs, uneven or low coverage and base errors. Interestingly, distributions of number of heteroplasmic variant calls are comparable for all whole-genome sequencing data sets, i.e. 1000G, HGDP and Pagani *et al*. (24) with a median number of heteroplasmic variant calls of 12, 14 and 12, respectively. Laricchia *et al*. (14) recently report for the gnomAD WGS data set of 56,434 individuals that most heteroplasmic variant calls are caused by NUMTs. This may also be the case here, because the Sudan mitochondrial amplicon sequencing data set has a much lower number of heteroplasmic variant calls (median 4), likely representing an upper bound of true heteroplasmic variants expected in blood of healthy individuals. However, the Egyptian mitochondrial amplicon sequencing data set has most heteroplasmic variant calls (median of 45). Finally, the Egyptian mummy data set has the widest range. Large numbers of heteroplasmic variant calls in mitochondrial sequencing observed here are caused by low and uneven coverage in conjunction with base errors. These base errors are possibly also reflecting input DNA quality, which decreases with time and environmental exposure; an example for this is the ancient DNA from Egyptian mummies. Thus, for accurate heteroplasmic variant identification down to low heteroplasmy levels, deep sequencing at an even coverage will be needed.

### Distribution of Western Asian and Northeast African mitochondrial haplogroup diversity

The sequential accumulation of polymorphisms in the mtDNA sequence over time across generations has led to the emergence of geographically isolable variation profiles and haplogroups. As a result, geographic regions have characteristic distributions of mtDNA haplogroups. The analysis of 9,264 worldwide sequences (see Methods) yielded haplogroup distributions for distinct geographical regions as shown in Figure 4a.

**Figure 4:**
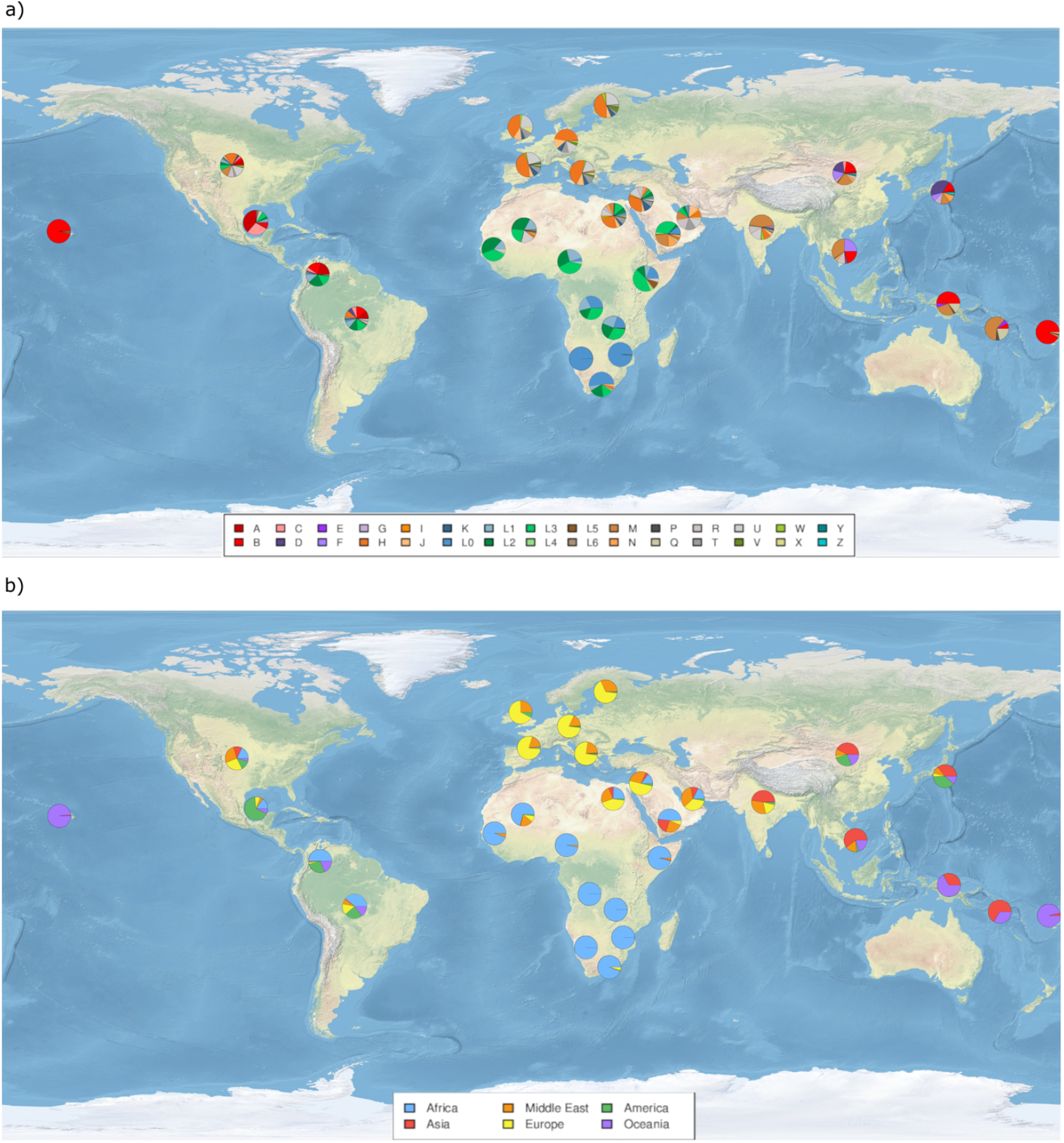
Worldwide haplogroup distributions by geographic region using top-level alphabetic clade denominators as well as first level subclades of haplogroup L. Please note that for some haplogroups geographic incidence of individual subclades can differ substantially (e.g. different B subclades occurring in America and Asia). a) Haplogroup level. b) Haplogroups have been assigned to continents of highest prevalence, representing their hypothetical regions of origin. We decided on the following assignment: Africa: L haplogroups; Asia: E, F, G, M, Y, Z; Middle East: N, R, U, I, W; Europe: H, J, K, T, V; America: A, C, D, X; Oceania: B, P, Q. Please note that tracing geographic origins of haplogroups is complex, e.g. due to autochthonous subclades, and the haplogroup levels and groupings we chose here are not representing all these relations, e.g. for haplogroup B.

As described previously, multiple haplogroups are predominant in different continental region. Here, we decided on the following assignment: Africa: L haplogroups; Asia: E, F, G, M, Y, Z; Middle East: N, R, U, I, W; Europe: H, J, K, T, V; Americas: A, C, D, X; Oceania: B, P, Q. Of note, we are aware that haplogroups overlap between continents, e.g., especially in Europe and the Middle East. The grouping above therefore does not entail exclusivity with respect to the listed haplogroups. Based on our assignment, we determined the relative proportion of haplogroups associated with different continents for each clustered group (see Figure 4b). Regions in Asia, North America, Northeast Africa, and the Middle East are particularly diverse. Historically, this can be attributed to events such as colonization and slave trade in the Americas as well as early human migration in the Middle East (39). In addition to its unique transcontinental location between Africa and Asia, Northeast Africa is adjacent to the center of modern human settlement in the world (15) and has been affected by complex events of prehistoric back migrations (40).

The relative frequency of mitochondrial haplogroups of the 641 QC filtered North and East African samples are depicted in Figure 5a and the individual haplogroup assignment per sample is given in Supplementary Table 11.

**Figure 5:**
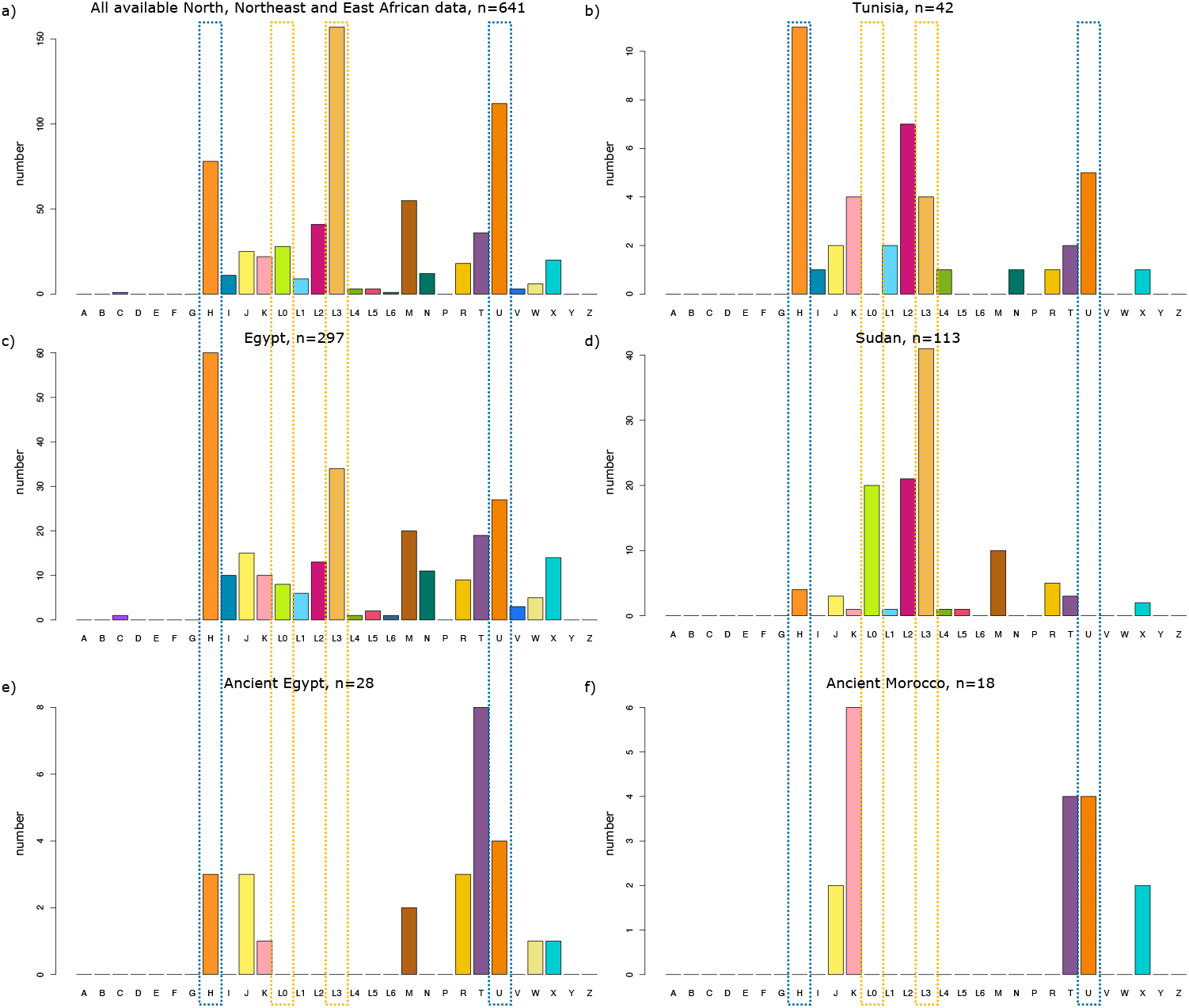
Histograms of haplogroup frequencies. Blue dashed line boxes indicate predominantly North African and yellow dashed line boxes predominantly East African haplogroups within this region. a) For all n=641 North, Northeast and East African samples passing the lenient QC filter and including Genbank sequences. This includes also Genbank sequences of studies addressing particular haplogroups (e.g. L3, M1 and U6) and thus is not a geographically representative distribution. b) Tunisian haplogroup numbers out of overall 42. c) Egyptian haplogroup numbers out of overall 297. d) Sudanese haplogroup numbers out of overall 113. e) Haplogroup numbers in ancient Egyptian mummies. f) Haplogroup numbers in ancient Moroccan individuals.

Country-specific haplogroup frequencies for Tunisia, Egypt, Sudan, ancient Morocco and ancient Egypt are shown in Figures 5b-f. Geographic neighborhoods are clearly reflected in the haplogroup distributions and indicate a geographic gradient from North Africa via Northeast Africa towards East Africa. This can be noted particularly for haplogroups H and U, that are most prevalent in Tunisia, less abundant in Egypt, and rare in Sudan. Conversely, L3 and L0 are prevalent in Sudan, and occur to a lesser extent in Egypt but are rarely observed in Tunisia. Interestingly, L2 is commonly observed in both Tunisia and Sudan, but rarer in Egypt. Overall, most haplogroup clades are observed in the Egyptian cohort (compiled from three data sets). This finding might be explained by the larger sample sizes and compilation of various data sets available for Egypt, compared to only one data set each for Tunisia and Sudan. Yet, more haplogroup clades reach high fractions compared to Tunisia and Sudan, supporting the hypothesis that Egypt is the region of highest mitochondrial DNA diversity within North- and East Africa.

### Northeast African mitochondrial lineages cover nearly all major worldwide clades

In order to elucidate the haplogroup abundance in North and East Africa relative to the rest of the world, we computed a phylogenetic tree from the 641 sequences in combination with the 1000G and HGDP reference data. The resulting trees in Figures 6 and 7 demonstrate that nearly all worldwide mitochondrial clades are represented in North and East Africa and, apart from the haplogroup M that is mainly represented by M1, a variety of subclades occurs. This is a haplogroup known to be restricted to North and East Africa that is absent from the 1000G and present in HGDP data only once.

**Figure 6:**
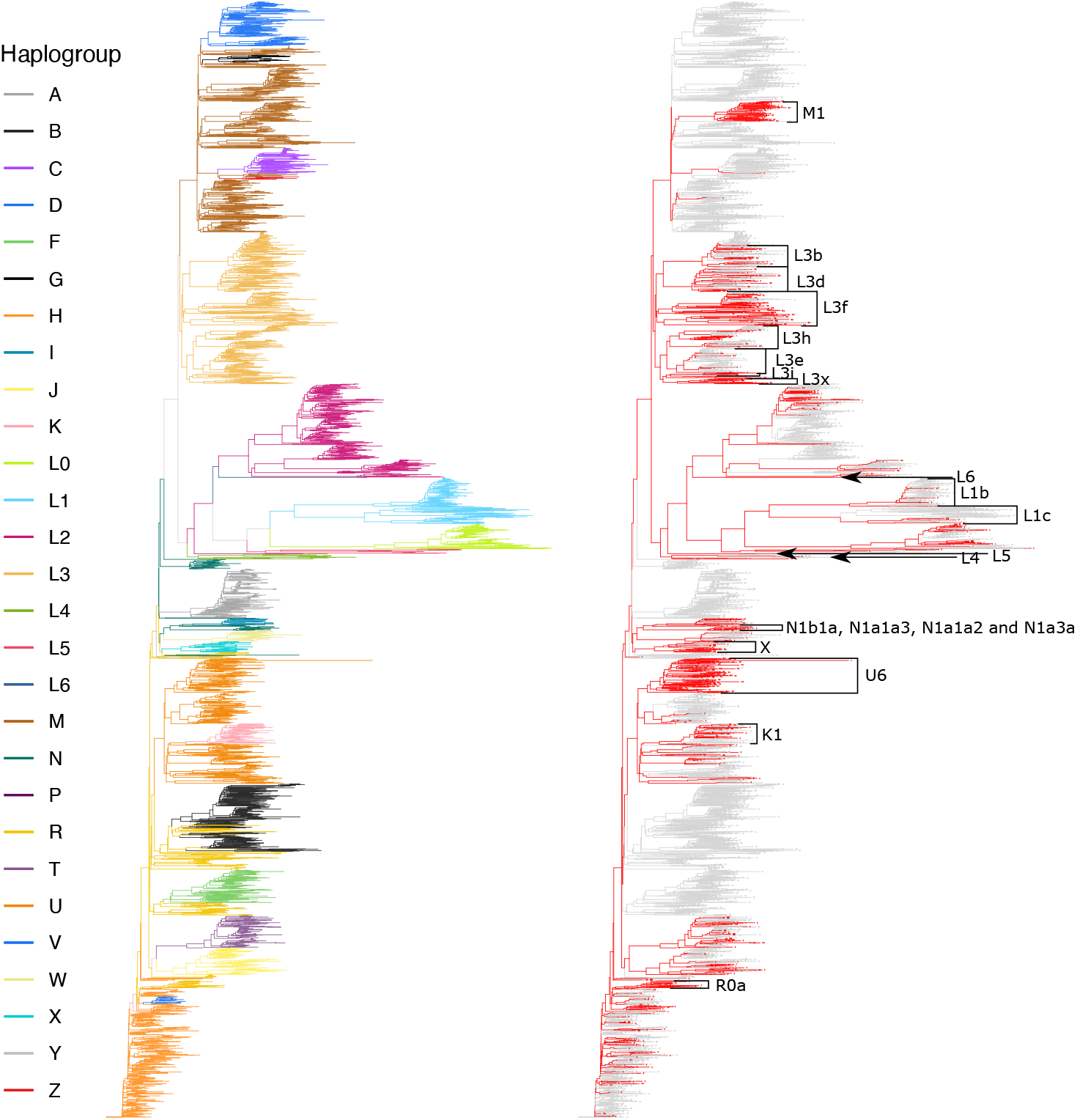
Phylogenetic tree of 641 North and East African mitochondrial sequences together with sequences from 1000G as worldwide reference data sets. This tree was computed by a Neighbor Joining-based tree and is colored according to haplogroup assigned by haplocheck (left) and colored according to data source, with North, Northeast and East African mitochondrial sequences compiled in this study shown in red and reference data shown in gray (right).

**Figure 7:**
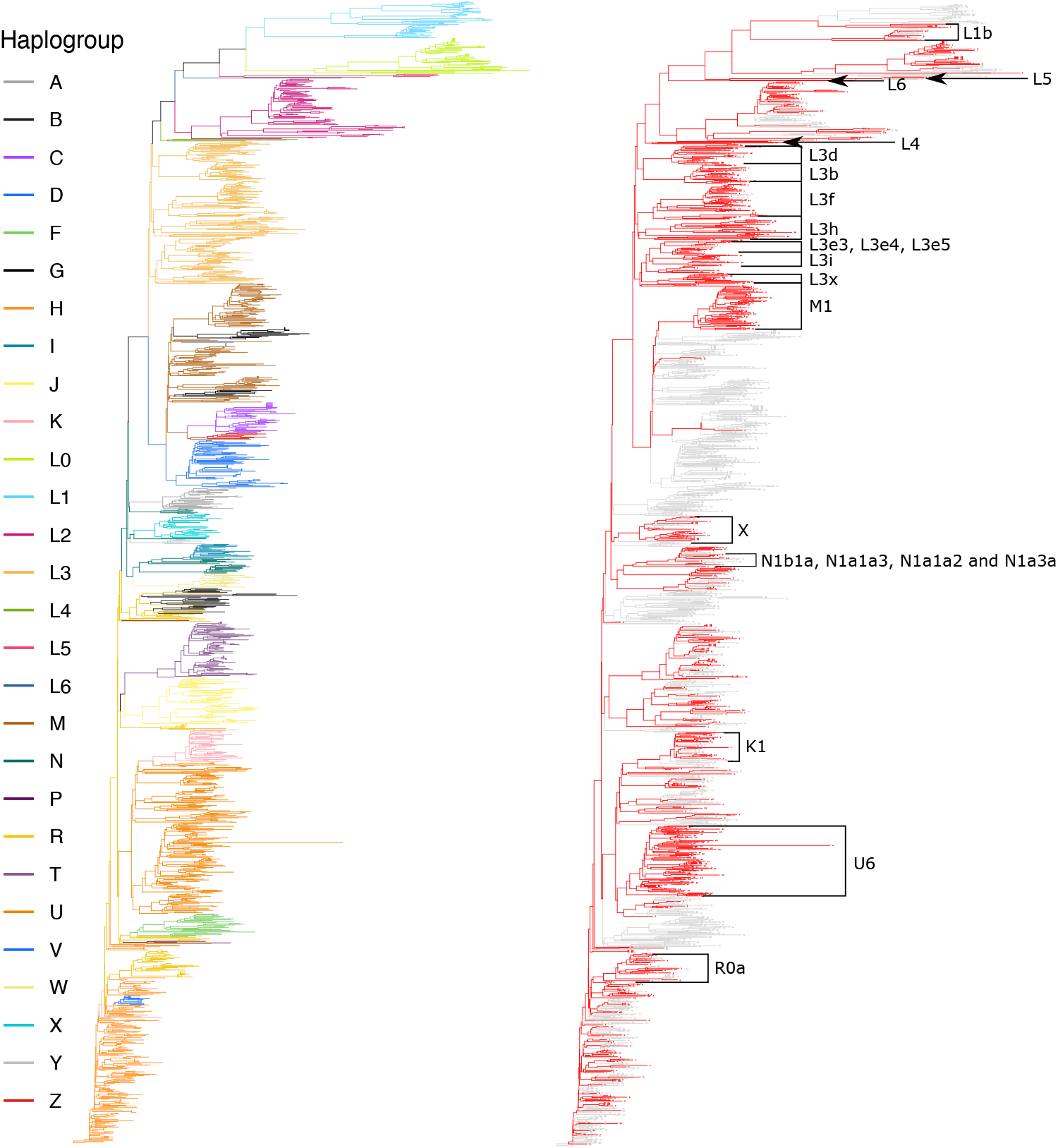
Phylogenetic tree of 641 North and East African mitochondrial sequences together with sequences from HGDP as worldwide reference data sets. This tree was computed by a Neighbor Joining-based tree and is colored according to haplogroup assigned by haplocheck (left) and colored according to data source, with North, Northeast and East African mitochondrial sequences compiled in this study shown in red and reference data shown in gray (right).

The worldwide haplogroups not observed in North and East Africa are A, B, D, E, F, G, P, Y and Z, which are predominantly North Asian, Northeast Asian, East Asian, Southeast Asian, American or Oceanian.

The predominantly European haplogroup H is more common in Tunisia and Egypt and less common in Sudan. The same applies to the haplogroups I, J, K, N and U.

Haplogroup I is a sister clade of N1a1b, as described earlier (41). Haplogroup J is also observed in both ancient data sets. North African sequences belong to J1 as well as J2 and are with respect to RSRS more deep-rooting than 1000G and HGDP sequences.

Haplogroup K represents a subgroup of U8, as expected (42). In the North and East African data, only K1 was observed, whereas 1000G and HGDP samples also covered K2.

African haplogroups L0, L1, L2 and L3 show a high abundance in Egypt, Sudan and Tunisia with the exceptions of L0 in Tunisia and L1 in Sudan being absent and lowly abundant, respectively. L3 is the founder of haplogroups N and M, which have given rise to all worldwide haplogroups outside of Africa, and thus L3 has been studied extensively and exclusively (33,40). Haplogroup L3 is observed in all data sets, but predominantly in Sudan with a share of 40%. This is in line with previous studies that suggest East Africa as the origin of L3 (33).

Besides pan-African haplogroups, we observe haplogroups L4, L5 and L6. All these haplogroups have previously been observed mainly in East Africa (43,44). Due to low prevalence and due to being restricted to this region, they have not been studied by explicitly sequencing mtDNA sequences of these haplogroups.

All but one sample relating to North and East African M haplogroups belong to the M1 clade, which is the only M subclade in Africa (45). The data sets compiled here cover a large diversity of M1. Haplogroup M1 has neither been observed in the comparably small Tunisian nor in the ancient Moroccan data set, but was present in the ancient Egyptian mummy data.

Sequences belonging to the N subhaplogroups build subclades that are located at different positions in the phylogenetic trees. All of them occur in the same phylogenetic branch, the same holds true for the haplogroup R. In the R subclade, we only observe haplogroups R0a (n=17) in North and East Africa, which previously has been described to be worldwide most frequent in the Middle East and North and East Africa (46), and encounter haplogroup R among the filtered sequences of an ancient Egyptian mummy once.

Nearly all T subclades are detected in North or East Africa, and their proportion is elevated in both Moroccan and Egyptian ancient data sets. The presence of haplogroup T, as well as its sister clade J, has previously been observed and considered to be of Near Eastern origin (25,36).

North and East African haplogroup U mtDNAs cover all subclades (U1 to U8) except the rare U9 clade. Haplogroup U6 has been studied in a targeted way (30,31,34) previously. Together with M1, it is the only clade of a worldwide haplogroup that is autochthonous in Africa, i.e., predominantly occurring there. Corresponding U6 sequences from (30) and (31) integrated into our data compilation as well as Egyptian and worldwide reference data support this notion.

Finally, haplogroup X is present in data from all North and East African regions, including data from three ancient individuals. The North and East African dataset contains 12 mtDNA sequences belonging to the phylogenetically old subclade X1, which is largely restricted to North Africa (47). The coalescence of X in the phylogenetic tree with 1000G and with HGDP, which both do not contain X1 mtDNA sequences, differs between both references and from the PhyloTree reference, involving N9b and R11 as sister clades, both of which are haplogroups that are poorly represented in the 1000G and HGDP reference data. In order to accurately resolve relationships between haplogroup X and other clades, more mtDNA sequences will be needed.

### Clades occurring predominantly in North and East Africa are often poorly characterized

We next investigated North and East African haplogroup profiles within the geographic context using selected 1000G and HGDP populations from neighboring regions (Figure 8). A corresponding histogram is shown in Supplementary Figure 1.

**Figure 8:**
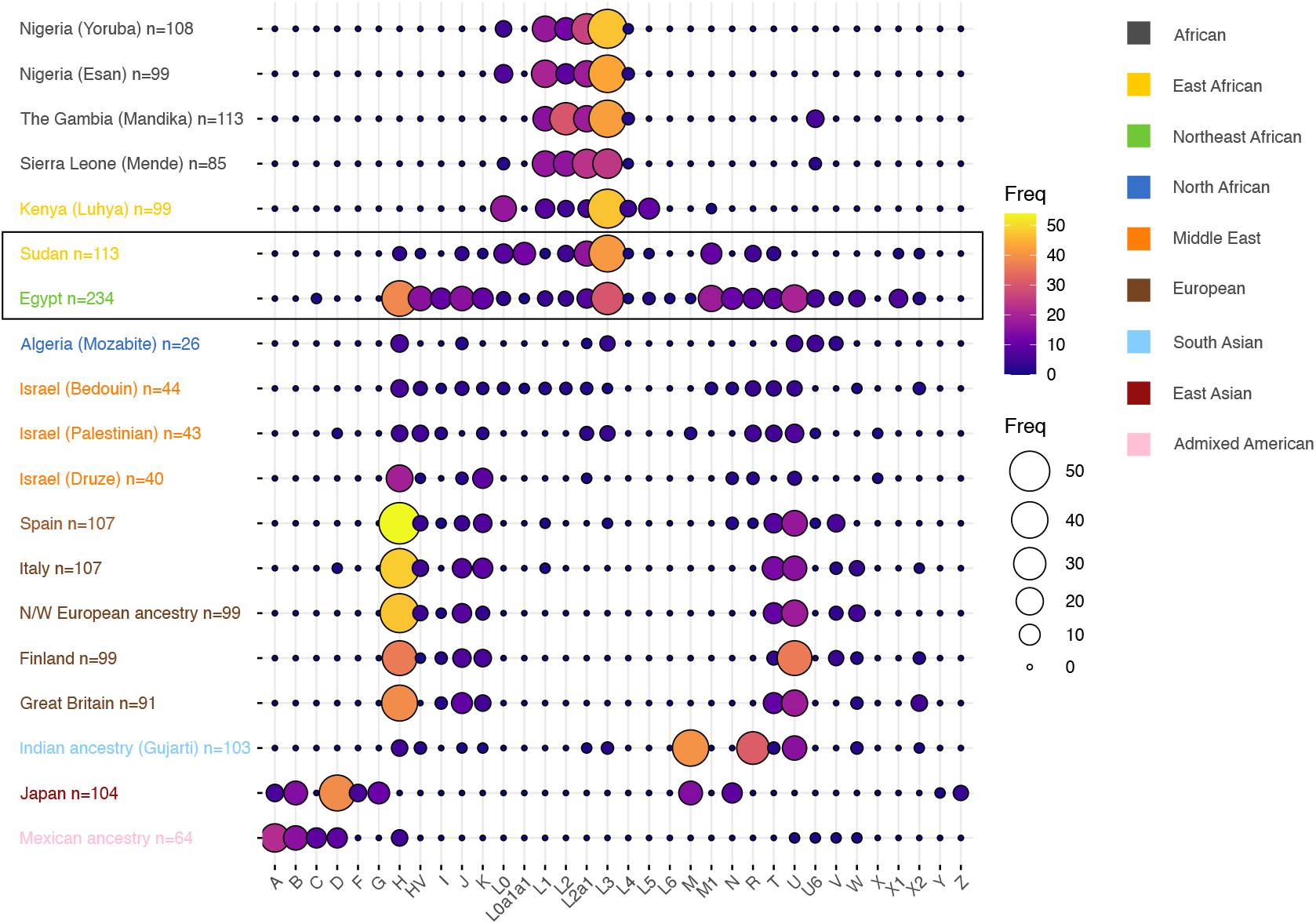
Haplogroup frequencies of selected 1000G and HGDP populations together with the Sudanese and Egyptian haplogroups of this work (highlighted in box). The populations are colored according to geographic region. Shown in the histogram are clades that are particularly relevant and/or prevalent in North or East Africa and discussed in this work., e.g. L0a1a1 and L2a1. Note that a sequence is counted only once, i.e., a sequence of haplogroup M1 is not counted towards haplogroup M.

This diagram shows that both Egyptian and Sudanese mitochondrial profiles are mediating between the profiles of populations assessed by these public references, constituting a genetic continuum between them. For example, 1000G covers African, European, South Asian, East Asian and Admixed American populations, none of which represents mitochondrial haplogroup occurrences observed in Sudan or Egypt according to data generated and compiled in this work (cf. Figure 8). Middle Eastern and North African populations of the HGDP reference show a combination of predominantly European, African and Asian haplogroups comparable to what is seen in Sudan and Egypt. However, HGDP population sample sizes are low and thus corresponding haplogroup distributions may not be representative.

Further, some haplogroups from the North and East African region are absent from both major global references 1000G and HGDP. This applies especially to specific haplogroups that exclusively or predominantly occur in North and/or East Africa, e.g. subclades of L0a1a1, L3, L4, L6, M1, U6 and X1. Some of them, with a particular interest to worldwide human prehistoric migration routes, have explicitly been addressed by studies and selectively been sequenced before, e.g., L3, M1 and U6 and are included in our study. However, local clades possibly not related to archaeogenetic hypothesis are more poorly characterized, e.g., L0a1a1. Finally, there is little information on rare haplogroups such as L6. They differ a lot from all other, well characterized haplogroups and as such may convey different mitochondrial function and disease susceptibility. Thus, also and especially such rare haplogroups will need better characterization in the context of mitochondrial genetics-based precision medicine.

### Variation suggesting novel haplogroups

To obtain a dataset of exceptional high-quality, we next selected (n=146) samples with high sequencing coverage and without predicted contamination (provided in Suppl. Table 12). The phylogenetic tree of these sequences with corresponding haplogroups is depicted in Figure 9.

**Figure 9:**
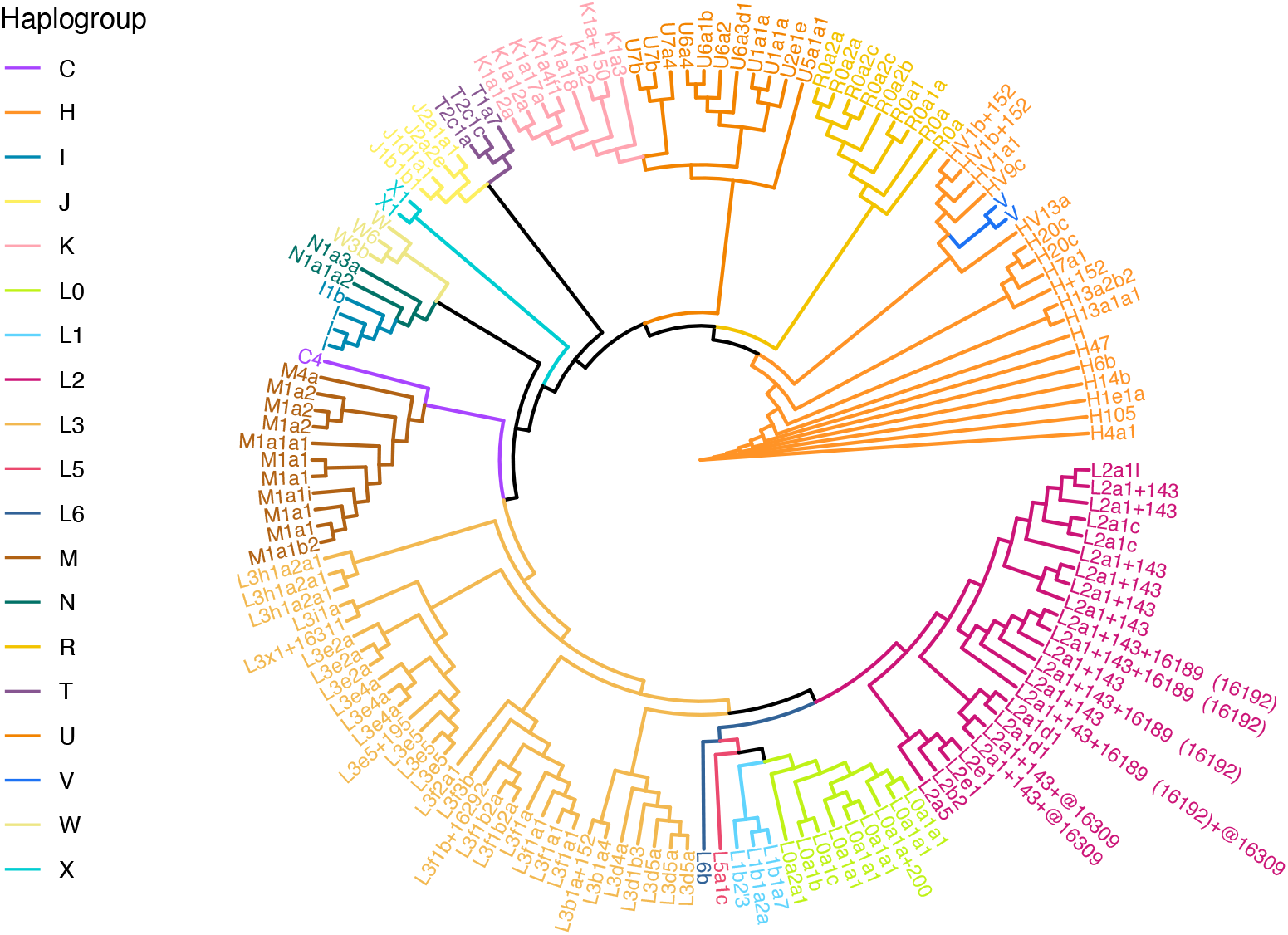
Phylogenetic tree of the 146 exceptional high-quality sequences used for detection of novel haplogroups. This tree was computed by a Neighbor Joining-based tree. The circular tree depicts the topological information, while branch lengths are not preserved. An alternative visualization preserving branch lengths is shown in Supplementary Figure 2.

In the search for novel and previously undefined subclades, we investigated this data set using the phantom mutation module of the online haplogrep2 tool. In this module, sequences are checked for recurrent variants that are not haplogroup-defining and that have a ‘Soares score’ of less than or equal to 2, i.e., having been detected no more than twice in a large collection of mitochondrial sequences (according to Table S3 of Soares et al. (48)). We checked for such variants occurring in samples of the same haplogroup to ensure that they were likely specific to haplogroup carriers and defined a novel, previously undefined subclade. A summary is provided in Table 1.

For ten putatively novel haplogroups, we find further support among samples with lower sequencing coverage. In seven cases, we find one novel, recurrent variant in at least two individuals of the same haplogroup, for example variant 8017T in overall 12 carriers of L0a1a1. In eight cases, more than one recurrent variant was detected within carriers of the same haplogroup, e.g. six carriers of L3f1a1 share six novel variants (12441C; 736T; 328G;15151G; 13967T; 14554G). Thereby, 14 of 30 potential novel haplogroup variants have a Soares score of 0 (47%), 9 have a score of 1 (30%) and the remaining 7 have a score of 2 (23%). The haplogroups of which potential novel subhaplogroups have been identified are either haplogroups that are largely restricted to North and/or East Africa, e.g., L2a1, M1, and different L3 clades (d, e and f), or are potential local clades diverging from basic haplogroups, e.g., I and V.

### Variation suggesting revision of phylogenetic relationships

For the 146 exceptional high-quality sequences, we examined the phylogenetic tree according to the PhyloTree reference to search for potential new haplogroups using a function from the web-based haplogrep2 tool (see Supplementary Figure 3). We manually investigated all variants present in the mtDNA sequence as well as all variants expected for the haplogroup assigned, but found to be missing in the sequence. We found two clades, L0a1a1 and L2a1, which were better represented, when revising or extending the phylogenetic tree provided by PhyloTree (Figure 10).

**Figure 10:**
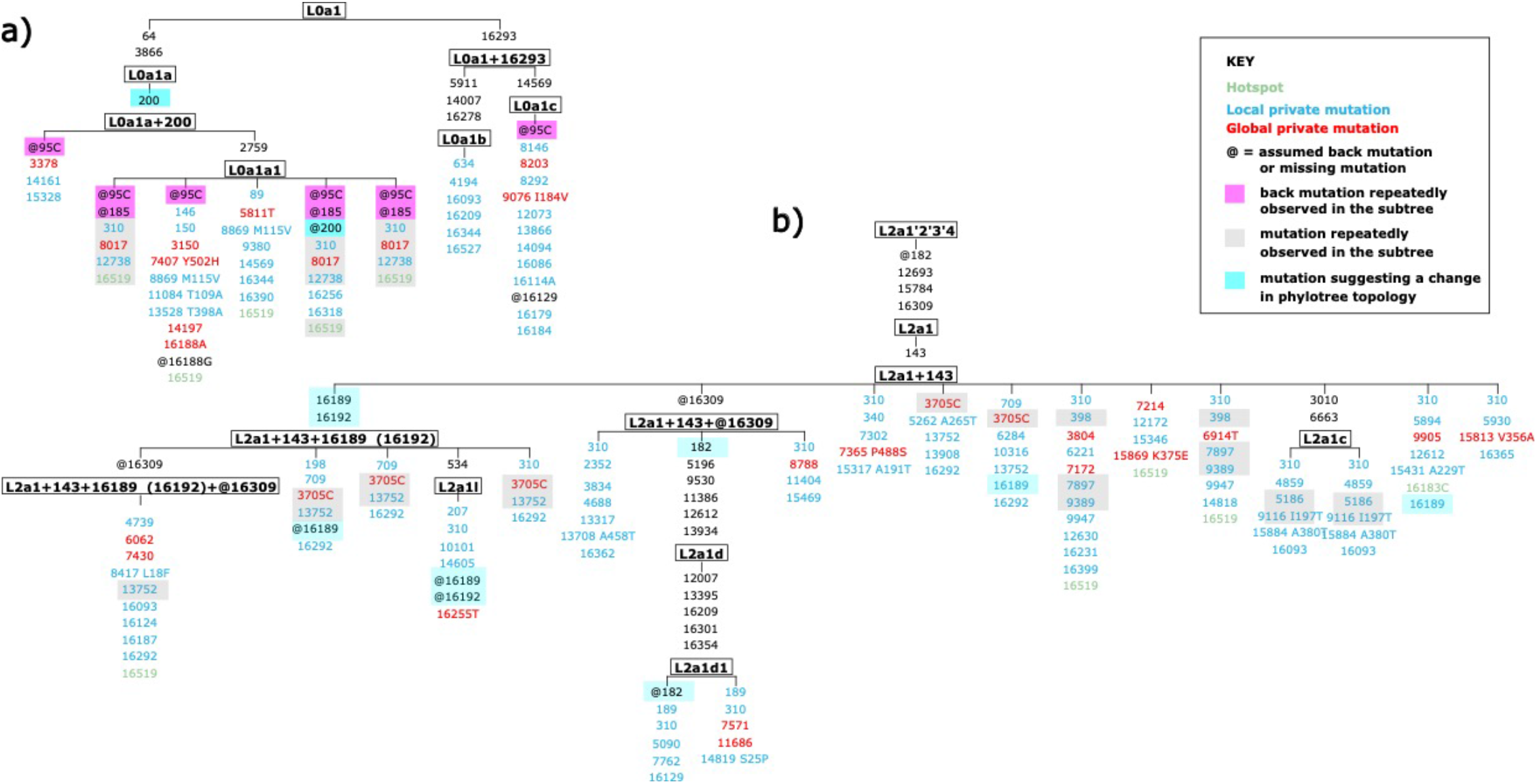
Phylogenetic tree according to the PhyloTree reference for those samples that were assigned to subclades a) L0a1 and b) L2a1. We highlight with magenta, gray and turquoise boxes those variants that indicate that a different phylogenetic tree may better explain the samples from North and East Africa, see legend in Figure a). Variants denoted for the leaves of the tree, i.e. samples, denote variant differences w.r.t. the haplogroup assigned. Shared back mutations (magenta) denote that parental haplotypes may be missing in PhyloTree. Mutations repeatedly observed in a subtree (gray) suggest that child haplogroups are missing. Variants that occur in multiple samples and differ from variants defining a parental haplogroup (turquoise) suggest that different variant combinations and haplogroup specifications may better explain North and East African mtDNA sequences.

There are eight samples assigned to L0a1 sub-haplogroups. In most of them, the haplogroup-defining variant 95C is not observed, which is already noted within the PhyloTree reference. Further, three samples show a back mutation at position 185. Such a back mutation is expected for haplogroup L0a2 according to PhyloTree, so meaning the samples we sequenced may link clades L0a1 and L0a2. In one sample, an expected mutation at position 200 was not observed, showing the possibility that 95C and 185 together with 310, 8017, 12738 and possibly 16519 are a hitherto undescribed subclade of L0a1.

The second branch of interest is L2a1. Out of 19 samples to which sub-haplogroups of L2a1 have been assigned, most showed extensive, recurrent deviations from PhyloTree (Figure 10 b). For example, resolving the haplogroup variants 16189 and 16192 within the PhyloTree reference seems not representative for North and East African samples. Further, variants 3705C, 13752, 7897, 9389, 5186 and 9116, which are recurrent in the L2a1 clade, are not represented in sequences appearing in PhyloTree, yet. However, if included, they would implicate changes in phylogeny.

## Conclusion

We investigated whether current global mitochondrial genetic data adequately represents the genetic mitochondrial variation of the North and East African regions. Genetic data from this region are sparse until now, despite an apparent interest due to autochthonous ancestry components being identified (22,49) and despite the importance of the region as origin or transit of modern humans outside of Africa. Besides an in-depth analysis of mitochondrial sequence data underlying Egyptian haplogroup distributions reported previously (22), we have generated novel sequence data of mitochondrial genomes from Sudan, another region with few genetic data available. We characterized North and East African mitochondrial DNA variation by combining these data with publicly available sequence data as well as with mitochondrial DNA sequences from Genbank relating to 10 other studies. This work has resulted, to the best of our knowledge, in the largest mitochondrial data collection from this region to date.

With a stringent quality control via the novel tool haplocheck, we related North and East African phylogenetics to the worldwide context by restricting the analysis to mitochondrial sequences unaffected by potential contamination or other shortcomings such as insufficient sequencing coverage. An additional filter for a minimal 1000x sequencing coverage results in exceptionally high-quality mitochondrial sequences, which makes us very confident that novel mitochondrial variations, novel haplogroups and observed PhyloTree inconsistencies are indeed caused by a hitherto insufficient genetic representation of this geographic region. Thus this work complements recent efforts to improve mitochondrial reference data (50,51).

We find that North and East African mitochondrial genomes are phylogenetically diverse and cover many of the major haplogroups seen in global populations. Likewise, specific subclades are prevalent in the region, supporting that North and East Africa, and particularly Northeast Africa harbors ancestry components typically attributed to Africa, Europe and Asia (22), all of which are likely present in the region since prehistoric times (52). Although our study focused on assessing the state of mtDNA reference data for precision medicine, the combined data allowed to rediscover geographic differences in genetic ancestry that many earlier studies related to prehistorical and historical conditions and events (53), e.g. migration between Egypt and Sudan along the Nile river (54). Further, several North or East African clade representatives highlighted by our work and not studied in-depth so far may still add further information to worldwide maternal human genetic history (e.g., the haplogroups L4, L6, N, I and X).

In conclusion, the initial North and East African mitochondrial reference dataset compiled as part of our work calls for further extension in order to support genetics-based precision medicine.

## Material and Methods

### Haplogroup data compilation for worldwide mitochondrial haplogroup distribution

Based on a comprehensive literature research, n=7,414 publicly available mitochondrial (mt)DNA sequences with available haplogroup and geographic information in the form of the country of origin were collected. Additionally, data from two reference datasets, the 1000 Genomes dataset (19) (n=2,504) and the Human Genome Diversity Project (28) (n=828) were added, resulting in a total number of n=10,746 entries. As quality assurance, for all database entries haplogroups were reassigned using for mitochondrial sequences haplogrep2 (10) and for next generation sequencing-based cohorts haplocheck (26). Subsequently, entries were filtered such that only those remained that were representative for the population of a region. Towards this, studies analyzing only specific haplogroups as well as data from archaeological studies was removed. Eventually, 9,264 entries were retained for further analysis.

### Objective geographic grouping for worldwide mitochondrial haplogroup distributions

For determining the relative amount of haplogroups per geographic region, we grouped the database entries into clusters of minimal geographic distance and having a size of at least 100 entries. At the beginning, only countries with less than 100 entries were considered, i.e., those that could not form a group on their own. Countries with an initial number less than 100 were assigned to group 1, all other countries to group 2. In order to classify a country from group 1, an attempt was made to combine it in a first step with other countries or already connected groups of countries in group 1. A continent-dependent maximum distance was set as a termination criterion. For Europe, this was set to 1,000 km, for Africa to 1,750 km, for the Middle East to 1,600 km, for Asia to 1,900 km, for the Americas to 2,600 km and for Oceania to 1,600 km. This distance was determined as a function of the average area of a country on the continent and the existing sample density. If after this first step, the sample number of a country or a group was still less than 100, the countries with an initial sample number greater than or equal to 100 were included as possible connection partners. In this second step, the country or group to be classified was in any case merged with the country from group 1 or group 2 with minimal distance.

### Mitochondrial DNA short-read sequencing

Sudanese samples were acquired under approval of the Central Institution Review Board, Al-Neelain University, Sudan, IRB Serial Number NU-IRB-17-7-7-91. All subjects gave written informed consent in accordance with the Declaration of Helsinki. Genomic DNA samples were processed for library preparation, using the Human mtDNA Genome protocol for Illumina Sequencing Platform (https://emea.support.illumina.com/content/dam/illumina-support/documents/documentation/chemistry_documentation/samplepreps_legacy/human-mtdna-genome-guide-15037958-01.pdf) as described previously (22,55) for Egyptian samples and 63 Sudanese samples. In brief, a primer set [MTL-F1 (AAAGCACATACCAAGGCCAC) and MTL-R1 (TTGGCTCTCCTTGCAAAGTT); MTL-F2 (TATCCGCCATCCCATACATT) and MTL-R2 (AATGTTGAGCCGTAGATGCC)] was used to amplify the mtDNA by long-range PCR (LR-PCR). Library preparation of the LR-PCR products was performed using a Nextera XT DNA Library Preparation Kit (Illumina Inc., CA, USA). For 96 Sudanese samples, we applied a slightly modified protocol to improve the evenness of the coverage. More specifically, the LR-PCR was conducted using another primer set [hMT-1_F (AAATCTTACCCCGCCTGTTT) and hMT-1_R (AATTAGGCTGTGGGTGGTTG); hMT-2_F (GCCATACTAGTCTTTGCCGC) and hMT-2_R (GGCAGGTCAATTTCACTGGT)] and the library preparation of the LR-PCR products of hMT1 and hMT2 primers was conducted using DNA Prep Tagmentation (Illumina Inc.). This approach allows the fragment size of the tagmented LR-PCR products to be evenly 300 bp, as the transposome reaction occurs every 300 bp on the beads, in contrast to the Nextera XT DNA library preparation Kit, by which transposome reaction occurs randomly, thus, results in various size of the tagmented LR-PCR products (i.e., larger fragments were removed by beads purification steps). The libraries prepared by both methods were further purified, concentration quantified using Qubit Fluorometer (Thermo Fisher GmbH, Dreieich, Germany) and the library size was determined by 2100 Bioanalyzer Instrument (Agilent, Santa Clara, CA, USA). The final library was sequenced on the Illumina MiSeq sequencing platform, using v2 chemistry (2 × 150 bp paired-end reads) (Illumina Inc.).

### Mitochondrial DNA long-read sequencing

Genomic DNA (gDNA) from peripheral blood for the long-read sequencing was prepared using Qiagen MagAttract HMW DNA kit (Qiagen, Hilden, Germany). Two GridION sequencing runs were conducted using one GridION Flow Cell version R10.3 (FLO-MIN111; Oxford Nanopore Technologies, Oxford, UK). Full-length mitochondrial DNA (16,595 bp) was amplified from gDNA by long-range PCR using a set of primers tagged with universal sequences (show in Italic): [US-hmt_F-2120 (*TTTCTGTTGGTGCTGATATTGC*GGACACTAGGAAAAAACCTTGTAGAGAGAG and US-hmt_R-2110 (*ACTTGCCTGTCGCTCTATCTTC*AAAGAGCTGTTCCTCTTTGGACTAACA)]. Fifty µl PCR reaction was prepared with 200 ng template gDNA, 50 nM of each forward and reverse primers, 25 µl LongAmp Hot Start Taq 2’ Master Mix (New England BioLabs, Frankfurt am Main, Germany). The PCR program was 94°C for 1 min; 30 cycles of 98°C for 10 s, and 68°C for 16 min; 72°C for 10 min; hold at 4°C. The PCR products were purified using GeneJET PCR purification Kit (Thermo Fisher Scientific GmbH, Dreireich, Germany). The purified PCR product were barcoded by PCR using the primer sets, which were the same sequences as Nanopore PCR Barcoding Expansion 1-96 (EXP-PBC096; Oxford Nanopore Technologies, UK), but were commercially synthesized (Biomers.net, Germany). The barcoding PCR condition was 95°C for 3 min; 15 cycles of 95°C for 15 sec, 62°C for 15 sec, and 65°C for 16 min; 65°C for 16 min; hold at 4°C.

The barcoded PCR products were further prepared for the DNA library using the Ligation Sequencing Kit (SQK-LSK110; Oxford Nanopore, UK) according to the manufacturer’s instructions. In brief, the end-repair, dA-tailing, sequencing adapter ligation, and final purification of DNA libraries were performed according to the manufacturer’s instructions using NEBNext Companion Module for Oxford Nanopore Technologies Ligation Sequencing (New England BioLabs GmbH, Germany). The GridION Flow Cell was primed and the DNA library was loaded according to the manufacturer’s instructions. Both sequencing durations were approximately 24 hours.

### Analysis of long-read MT sequencing data

Basecalling was performed with guppy version 6.0.1+652ffd1 and flowcell option FLO-MIN111, sequencing kit option SQK-LSK110 and barcode kit option EXP-PBC096, which defaults to guppy config file dna_r10.3_450bps_hac.cfg and model file template_r10.3_450bps_hac.jsn, i.e. using high accuracy base calling. Reads with a Q-score larger 10 were considered pass reads and kept for further analysis. PycoQC (56) was run for quality control and mapping performed with minimap2 against GRCh38, which includes rCRS. Haplogroups were assigned based on this BAM file using the haplocheck tool within mitoVerse (https://mitoverse.i-med.ac.at).

### Sudanese and Egyptian mitochondrial sequencing characteristics and QC

Running FastQC on all Sudanese and Egyptian mitochondrial sequencing FASTQ files resulted in no systematic, data quality compromising warnings or errors. Sudanese sequencing data has an average length of 140 bases, Egyptian sequencing comprises samples sequenced with 150 and samples sequenced with 300 bp target read length, resulting in an average length of 205 bases across the cohort. Average bases mapped per sample are about 18 Mb for the Sudanese cohort and about 30 Mb for the Egyptian cohort, amounting to an approximate average coverage of more than 1000X. In the Sudanese cohort, five samples have less than 1000 reads and are considered dropouts. The Egyptian cohort does not contain samples with less than 1000 reads.

### Variant calling, haplogroup assignment and contamination detection

Raw reads from FASTQ files of Sudanese and Egyptian sequencing data were mapped to mitochondrial sequence MT of GRCh38, which resembles the historic revised Cambridge Reference Sequence (rCRS), a sequence of haplogroup H2a2a1. For samples with available BAM files, we used these directly. Our analysis is largely based on haplocheck (version 1.1.2) (26) and its internal accompanying tools haplogrep2 (10) and mutserve (version 1.3.4) as implemented via the mtDNA-server (57). Variant caller mutserv was run with default settings, which amounts to reporting heteroplasmic variants at a level larger than 1% (--level 0.01), calling deletions (--deletions), base alignment quality turned off (--noBaq), considering a mapping quality of 20 (--mapQ 20) and a base quality of 20 (--basQ).

### Phylogenetic analyses

In order to obtain a FASTA format mitochondrial sequence that relates to the haplogrep2 output reported by haplocheck, we perform variant calling with mutserv using the default parameters also used within haplocheck. Then we select from the VCF those variants for which the major base is the non-reference base. Using bcftools consensus, we then convert the VCF file into FASTA. Note that mutserv does not call insertions, only deletions which are denoted with a gap symbol (within FASTA, it is “-“), thus ensuring that the sequence length is always exactly the length of rCRS, i.e. 16,569 bases, and that sequences align correctly without re-alignment. The same holds for the Genbank files processed by McInerny et al. (29). Thus, we can simply combine FASTA files of cohorts by pasting them into one combined FASTA file. The combined FASTA file was then loaded into Jalview (58) where a Neighbour Joining-based tree was computed and exported into Newick format. This tree was then visualized with ggtree (59) using various graphical settings.

## Supporting information

Supplement with Suppl. Figures 1, 2 and 3

## Ethics statement

This study was approved by the Central Institution Review Board, Al-Neelain University, Sudan, IRB Serial Number NU-IRB-17-7-7-91. All subjects gave written informed consent in accordance with the Declaration of Helsinki.

## Data availability

The public data compiled as well as the corresponding publications and repository IDs are provided in Supplementary Table 1. Sudanese raw sequencing data generated as part of this study is available from the European Genome Phenome Archive (EGA) under study ID EGAD00001008315. The analysis workflow used to run haplocheck, perform variant calling and generate and compile mitochondrial sequences is available at https://github.com/iwohlers/2020_mt_analyses.

## Acknowledgements

The authors thank all volunteers who donated blood for the study as well as everyone involved in the collection of blood samples. We also thank Miriam Freitag for excellent technical assistance. IW, HB and SI acknowledge funding by the Deutsche Forschungsgemeinschaft (DFG, German Research Foundation) under Germany’s Excellence Strategy—EXC 22167-390884018. AF, IS, HB and IW acknowledge support through the high-performance computer cluster (OMICS-Cluster) of the University of Lübeck.

## References

1. Wallace DC, Fan W, Procaccio V. Mitochondrial energetics and therapeutics. Annu Rev Pathol. 2010;5:297–348.

2. Anderson S, Bankier AT, Barrell BG, de Bruijn MH, Coulson AR, Drouin J, et al. Sequence and organization of the human mitochondrial genome. Nature. 1981 Apr 9;290(5806):457–65.

3. Chandel NS. Evolution of Mitochondria as Signaling Organelles. Cell Metab. 2015 Aug 4;22(2):204–6.

4. Prasun P. Mitochondrial dysfunction in metabolic syndrome. Biochim Biophys Acta Mol Basis Dis. 2020 Oct 1;1866(10):165838.

5. Wang W, Zhao F, Ma X, Perry G, Zhu X. Mitochondria dysfunction in the pathogenesis of Alzheimer’s disease: recent advances. Mol Neurodegener. 2020 May 29;15(1):30.

6. Hsu CC, Tseng LM, Lee HC. Role of mitochondrial dysfunction in cancer progression. Exp Biol Med (Maywood). 2016 Jun;241(12):1281–95.

7. Wallace DC. A mitochondrial paradigm of metabolic and degenerative diseases, aging, and cancer: a dawn for evolutionary medicine. Annu Rev Genet. 2005;39:359–407.

8. van Oven M, Kayser M. Updated comprehensive phylogenetic tree of global human mitochondrial DNA variation. Hum Mutat. 2009 Feb;30(2):E386–394.

9. van Oven M. PhyloTree Build 17: Growing the human mitochondrial DNA tree. Forensic Science International: Genetics Supplement Series. 2015 Dec 1;5:e392–4.

10. Weissensteiner H, Pacher D, Kloss-Brandstätter A, Forer L, Specht G, Bandelt HJ, et al. HaploGrep 2: mitochondrial haplogroup classification in the era of high-throughput sequencing. Nucleic Acids Res. 2016 08;44(W1):W58–63.

11. Rishishwar L, Jordan IK. Implications of human evolution and admixture for mitochondrial replacement therapy. BMC Genomics. 2017 Feb 8;18(1):140.

12. Lippold S, Xu H, Ko A, Li M, Renaud G, Butthof A, et al. Human paternal and maternal demographic histories: insights from high-resolution Y chromosome and mtDNA sequences. Investig Genet. 2014;5:13.

13. Rieux A, Eriksson A, Li M, Sobkowiak B, Weinert LA, Warmuth V, et al. Improved calibration of the human mitochondrial clock using ancient genomes. Mol Biol Evol. 2014 Oct;31(10):2780–92.

14. Laricchia KM, Lake NJ, Watts NA, Shand M, Haessly A, Gauthier L, et al. Mitochondrial DNA variation across 56,434 individuals in gnomAD. Genome Res. 2022 Mar;32(3):569– 82.

15. Bergström A, Stringer C, Hajdinjak M, Scerri EML, Skoglund P. Origins of modern human ancestry. Nature. 2021 Feb;590(7845):229–37.

16. Behar DM, van Oven M, Rosset S, Metspalu M, Loogväli EL, Silva NM, et al. A “Copernican” reassessment of the human mitochondrial DNA tree from its root. Am J Hum Genet. 2012 Apr 6;90(4):675–84.

17. Chinnery PF. Precision mitochondrial medicine. Cambridge Prisms: Precision Medicine. 2022 Nov 15;1–23.

18. Gorman GS, Schaefer AM, Ng Y, Gomez N, Blakely EL, Alston CL, et al. Prevalence of nuclear and mitochondrial DNA mutations related to adult mitochondrial disease. Ann Neurol. 2015 May;77(5):753–9.

19. 1000 Genomes Project Consortium, Auton A, Brooks LD, Durbin RM, Garrison EP, Kang HM, et al. A global reference for human genetic variation. Nature. 2015 Oct 1;526(7571):68–74.

20. Karczewski KJ, Francioli LC, Tiao G, Cummings BB, Alföldi J, Wang Q, et al. The mutational constraint spectrum quantified from variation in 141,456 humans. Nature. 2020;581(7809):434–43.

21. Taliun D, Harris DN, Kessler MD, Carlson J, Szpiech ZA, Torres R, et al. Sequencing of 53,831 diverse genomes from the NHLBI TOPMed Program. Nature. 2021 Feb;590(7845):290–9.

22. Wohlers I, Künstner A, Munz M, Olbrich M, Fähnrich A, Calonga-Solís V, et al. An integrated personal and population-based Egyptian genome reference. Nat Commun. 2020 Sep 18;11(1):4719.

23. El-Attar EA, Helmy Elkaffas RM, Aglan SA, Naga IS, Nabil A, Abdallah HY. Genomics in Egypt: Current Status and Future Aspects. Front Genet. 2022;13:797465.

24. Pagani L, Schiffels S, Gurdasani D, Danecek P, Scally A, Chen Y, et al. Tracing the route of modern humans out of Africa by using 225 human genome sequences from Ethiopians and Egyptians. Am J Hum Genet. 2015 Jun 4;96(6):986–91.

25. Schuenemann VJ, Peltzer A, Welte B, van Pelt WP, Molak M, Wang CC, et al. Ancient Egyptian mummy genomes suggest an increase of Sub-Saharan African ancestry in post-Roman periods. Nat Commun. 2017 May 30;8:15694.

26. Weissensteiner H, Forer L, Fendt L, Kheirkhah A, Salas A, Kronenberg F, et al. Contamination detection in sequencing studies using the mitochondrial phylogeny. Genome Res. 2021 Jan 15;

27. Brandstätter A, Sänger T, Lutz-Bonengel S, Parson W, Béraud-Colomb E, Wen B, et al. Phantom mutation hotspots in human mitochondrial DNA. Electrophoresis. 2005 Sep;26(18):3414–29.

28. Bergström A, McCarthy SA, Hui R, Almarri MA, Ayub Q, Danecek P, et al. Insights into human genetic variation and population history from 929 diverse genomes. Science. 2020 20;367(6484).

29. McInerney TW, Fulton-Howard B, Patterson C, Paliwal D, Jermiin LS, Patel HR, et al. A globally diverse reference alignment and panel for imputation of mitochondrial DNA variants. BMC Bioinformatics. 2021 Sep 1;22(1):417.

30. Maca-Meyer N, González AM, Pestano J, Flores C, Larruga JM, Cabrera VM. Mitochondrial DNA transit between West Asia and North Africa inferred from U6 phylogeography. BMC Genet. 2003 Oct 16;4:15.

31. Olivieri A, Achilli A, Pala M, Battaglia V, Fornarino S, Al-Zahery N, et al. The mtDNA legacy of the Levantine early Upper Palaeolithic in Africa. Science. 2006 Dec 15;314(5806):1767–70.

32. Harich N, Costa MD, Fernandes V, Kandil M, Pereira JB, Silva NM, et al. The trans-Saharan slave trade - clues from interpolation analyses and high-resolution characterization of mitochondrial DNA lineages. BMC Evol Biol. 2010 May 10;10:138.

33. Soares P, Alshamali F, Pereira JB, Fernandes V, Silva NM, Afonso C, et al. The Expansion of mtDNA Haplogroup L3 within and out of Africa. Mol Biol Evol. 2012 Mar;29(3):915– 27.

34. Pennarun E, Kivisild T, Metspalu E, Metspalu M, Reisberg T, Moisan JP, et al. Divorcing the Late Upper Palaeolithic demographic histories of mtDNA haplogroups M1 and U6 in Africa. BMC Evol Biol. 2012 Dec 3;12:234.

35. Fregel R, Méndez FL, Bokbot Y, Martín-Socas D, Camalich-Massieu MD, Santana J, et al. Ancient genomes from North Africa evidence prehistoric migrations to the Maghreb from both the Levant and Europe. Proc Natl Acad Sci U S A. 2018 Jun 26;115(26):6774– 9.

36. Kujanová M, Pereira L, Fernandes V, Pereira JB, Cerný V. Near eastern neolithic genetic input in a small oasis of the Egyptian Western Desert. Am J Phys Anthropol. 2009 Oct;140(2):336–46.

37. Costa MD, Cherni L, Fernandes V, Freitas F, Ammar El Gaaied AB, Pereira L. Data from complete mtDNA sequencing of Tunisian centenarians: testing haplogroup association and the “golden mean” to longevity. Mech Ageing Dev. 2009 Apr;130(4):222–6.

38. Lüth T, Schaake S, Grünewald A, May P, Trinh J, Weissensteiner H. Benchmarking Low-Frequency Variant Calling With Long-Read Data on Mitochondrial DNA. Front Genet. 2022;13:887644.

39. Rodriguez-Flores JL, Fakhro K, Agosto-Perez F, Ramstetter MD, Arbiza L, Vincent TL, et al. Indigenous Arabs are descendants of the earliest split from ancient Eurasian populations. Genome Res. 2016 Feb;26(2):151–62.

40. Cabrera VM, Marrero P, Abu-Amero KK, Larruga JM. Carriers of mitochondrial DNA macrohaplogroup L3 basal lineages migrated back to Africa from Asia around 70,000 years ago. BMC Evol Biol. 2018 Jun 19;18(1):98.

41. Olivieri A, Pala M, Gandini F, Hooshiar Kashani B, Perego UA, Woodward SR, et al. Mitogenomes from two uncommon haplogroups mark late glacial/postglacial expansions from the near east and neolithic dispersals within Europe. PLoS One. 2013;8(7):e70492.

42. González AM, García O, Larruga JM, Cabrera VM. The mitochondrial lineage U8a reveals a Paleolithic settlement in the Basque country. BMC Genomics. 2006 May 23;7:124.

43. Kivisild T, Reidla M, Metspalu E, Rosa A, Brehm A, Pennarun E, et al. Ethiopian mitochondrial DNA heritage: tracking gene flow across and around the gate of tears. Am J Hum Genet. 2004 Nov;75(5):752–70.

44. Tishkoff SA, Gonder MK, Henn BM, Mortensen H, Knight A, Gignoux C, et al. History of click-speaking populations of Africa inferred from mtDNA and Y chromosome genetic variation. Mol Biol Evol. 2007 Oct;24(10):2180–95.

45. Metspalu M, Kivisild T, Metspalu E, Parik J, Hudjashov G, Kaldma K, et al. Most of the extant mtDNA boundaries in South and Southwest Asia were likely shaped during the initial settlement of Eurasia by anatomically modern humans. BMC Genet. 2004 Aug 31;5:26.

46. Gandini F, Achilli A, Pala M, Bodner M, Brandini S, Huber G, et al. Mapping human dispersals into the Horn of Africa from Arabian Ice Age refugia using mitogenomes. Sci Rep. 2016 May 5;6:25472.

47. Reidla M, Kivisild T, Metspalu E, Kaldma K, Tambets K, Tolk HV, et al. Origin and diffusion of mtDNA haplogroup X. Am J Hum Genet. 2003 Nov;73(5):1178–90.

48. Soares P, Ermini L, Thomson N, Mormina M, Rito T, Röhl A, et al. Correcting for purifying selection: an improved human mitochondrial molecular clock. Am J Hum Genet. 2009 Jun;84(6):740–59.

49. Hodgson JA, Mulligan CJ, Al-Meeri A, Raaum RL. Early back-to-Africa migration into the Horn of Africa. PLoS Genet. 2014 Jun;10(6):e1004393.

50. Dür A, Huber N, Parson W. Fine-Tuning Phylogenetic Alignment and Haplogrouping of mtDNA Sequences. Int J Mol Sci. 2021 May 27;22(11):5747.

51. Bolze A, Mendez F, White S, Tanudjaja F, Isaksson M, Jiang R, et al. A catalog of homoplasmic and heteroplasmic mitochondrial DNA variants in humans. bioRxiv. 2020 Jan 1;798264.

52. Serra-Vidal G, Lucas-Sanchez M, Fadhlaoui-Zid K, Bekada A, Zalloua P, Comas D. Heterogeneity in Palaeolithic Population Continuity and Neolithic Expansion in North Africa. Curr Biol. 2019 Nov 18;29(22):3953-3959.e4.

53. Lucas-Sánchez M, Serradell JM, Comas D. Population history of North Africa based on modern and ancient genomes. Hum Mol Genet. 2021 Apr 26;30(R1):R17–23.

54. Krings M, Salem AE, Bauer K, Geisert H, Malek AK, Chaix L, et al. mtDNA analysis of Nile River Valley populations: A genetic corridor or a barrier to migration? Am J Hum Genet. 1999 Apr;64(4):1166–76.

55. Russlies J, Fähnrich A, Witte M, Yin J, Benoit S, Gläser R, et al. Polymorphisms in the Mitochondrial Genome Are Associated With Bullous Pemphigoid in Germans. Front Immunol. 2019;10:2200.

56. Leger A, Leonardi T. pycoQC, interactive quality control for Oxford Nanopore Sequencing. Journal of Open Source Software. 2019;4(34):1236.

57. Weissensteiner H, Forer L, Fuchsberger C, Schöpf B, Kloss-Brandstätter A, Specht G, et al. mtDNA-Server: next-generation sequencing data analysis of human mitochondrial DNA in the cloud. Nucleic Acids Res. 2016 Jul 8;44(W1):W64–69.

58. Waterhouse AM, Procter JB, Martin DMA, Clamp M, Barton GJ. Jalview Version 2--a multiple sequence alignment editor and analysis workbench. Bioinformatics. 2009 May 1;25(9):1189–91.

59. Yu G. Using ggtree to Visualize Data on Tree-Like Structures. Curr Protoc Bioinformatics. 2020 Mar;69(1):e96.

