## Supplement with Suppl. Figures 1, 2 and 3 for "North and East African mitochondrial genetic variation needs further characterization towards precision medicine"

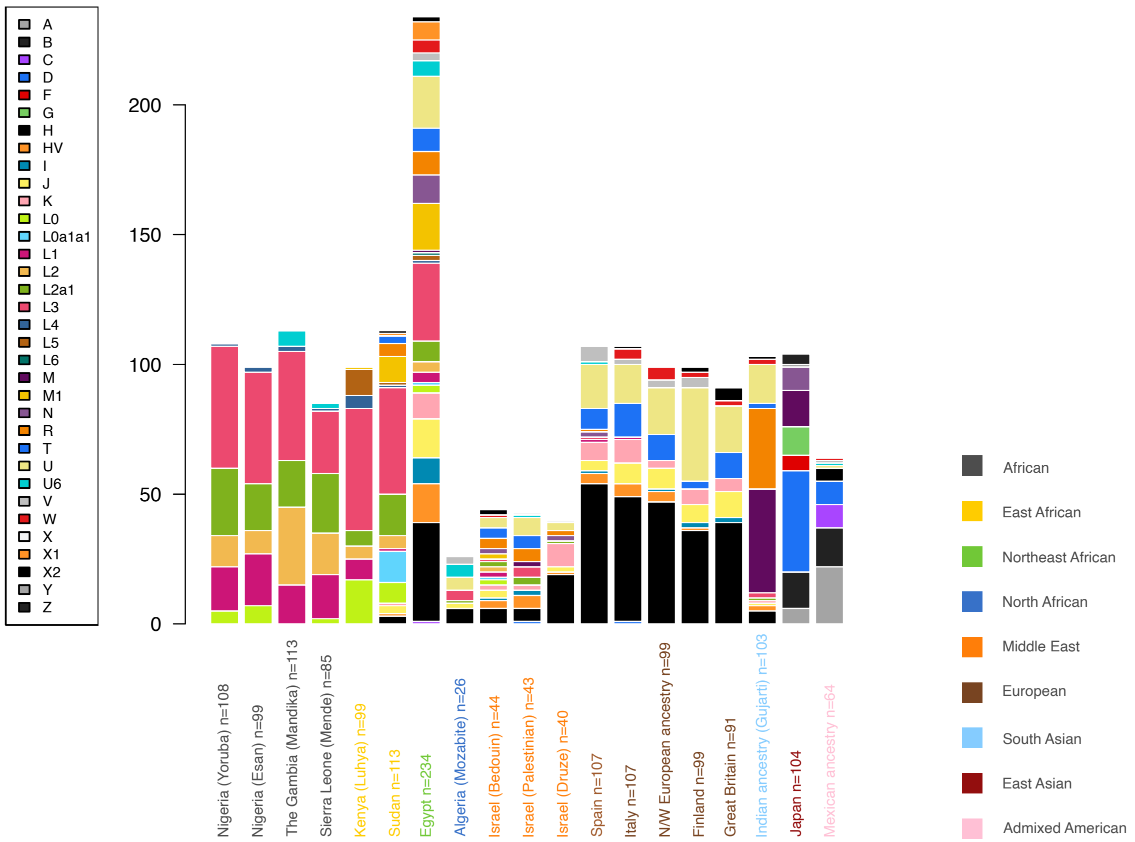

***Suppl. Figure 1:*** Histogram representation for Figure 6: Haplogroup frequencies of selected 1000G and HGDP populations together with the Sudanese and Egyptian haplogroups of this work. The populations are colored according to geographic region. Shown in the histogram are clades that are particularly relevant and/or prevalent in North or East Africa and discussed in this work., e.g. L0a1a1 and L2a1. Note that a sequence is counted only once, i.e., a sequence of haplogroup M1 is not counted towards haplogroup M.

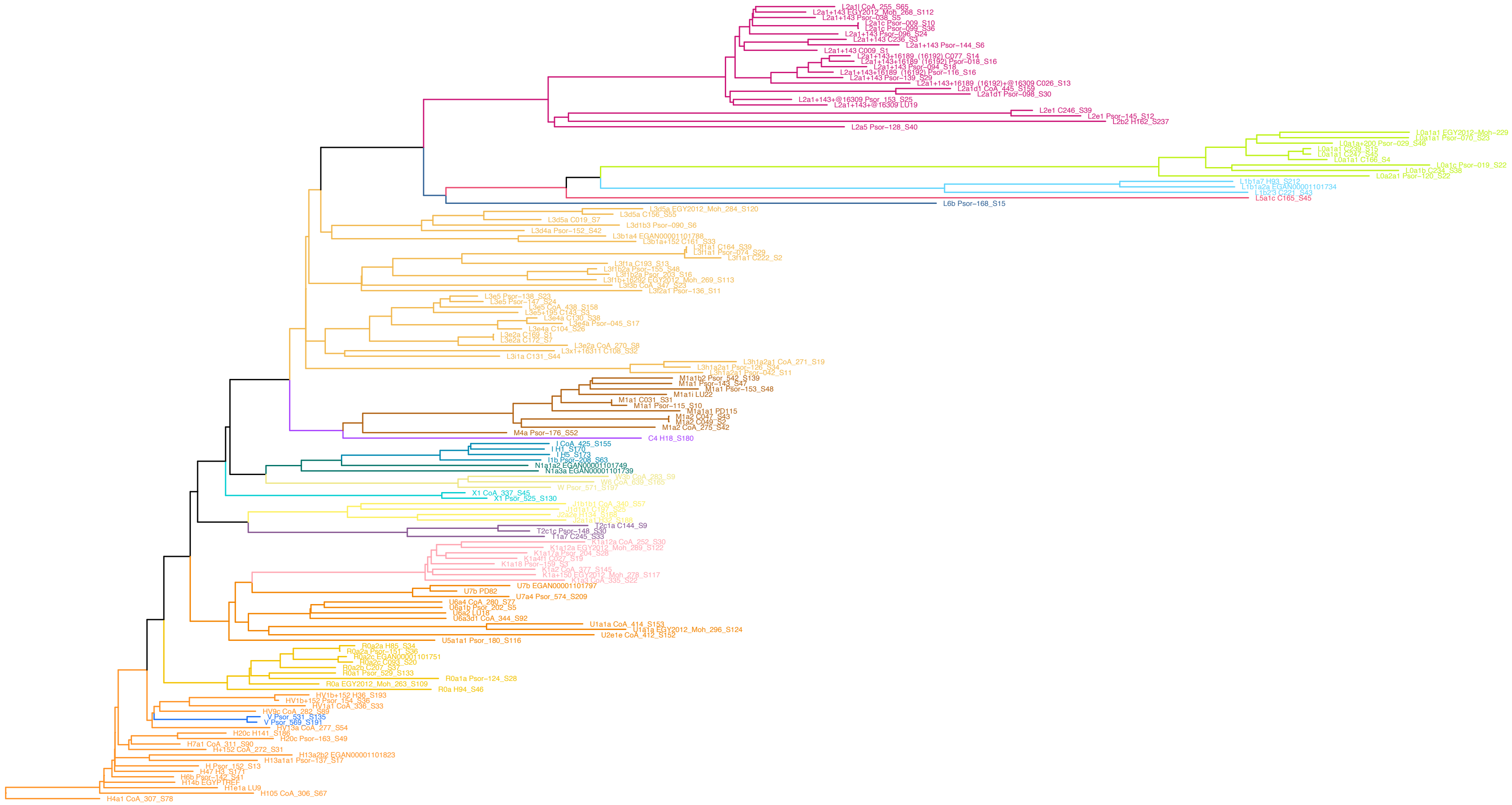

***Suppl. Figure 2:*** Phylogenetic neighbour joining-based tree of the n=146 exceptional high-quality sequences used for detection of novel haplogroups. Shown are two alternative representations of the tree in Figure 7, as well as the corresponding haplogroup histogram.

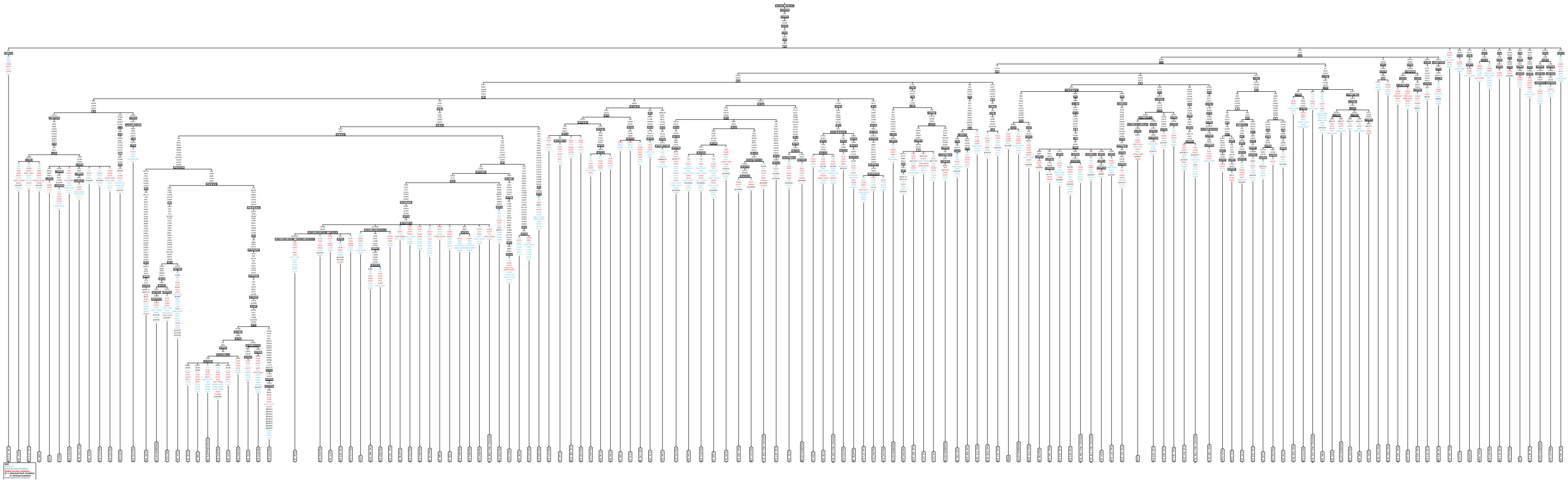

***Suppl. Figure 3:*** For the 146 exceptional high-quality sequences, the phylogenetic tree according to the PhyloTree reference, generated using a function from the web-based haplogrep2 tool.
